# Origin of wiring specificity in an olfactory map: dendrite targeting of projection neurons

**DOI:** 10.1101/2022.12.28.522173

**Authors:** Kenneth Kin Lam Wong, Tongchao Li, Tian-Ming Fu, Gaoxiang Liu, Cheng Lyu, Sayeh Kohani, Qijing Xie, David J Luginbuhl, Srigokul Upadhyayula, Eric Betzig, Liqun Luo

## Abstract

How does wiring specificity of neural maps emerge during development? Formation of the adult *Drosophila* olfactory glomerular map begins with patterning of projection neuron (PN) dendrites at the early pupal stage. To better understand the origin of wiring specificity of this map, we created genetic tools to systematically characterize dendrite patterning across development at PN type–specific resolution. We find that PNs use lineage and birth order combinatorially to build the initial dendritic map. Specifically, birth order directs dendrite targeting in rotating and binary manners for PNs of the anterodorsal and lateral lineages, respectively. Two-photon– and adaptive optical lattice light-sheet microscope–based time-lapse imaging reveals that PN dendrites initiate active targeting with direction-dependent branch stabilization on the timescale of seconds. Moreover, PNs that are used in both the larval and adult olfactory circuits prune their larval-specific dendrites and re-extend new dendrites simultaneously to facilitate timely olfactory map organization. Our work highlights the power and necessity of type-specific neuronal access and time-lapse imaging in identifying wiring mechanisms that underlie complex patterns of functional neural maps.

## INTRODUCTION

Organization of neuronal connectivity into spatial maps occurs widely in the nervous systems across species (Luo & Flanagan, 2007; Cang & Feldheim, 2013; Luo, 2021). For example, in the retinotopic map of the visual system, neighboring neurons in the input field project axons to neighboring neurons in the target field (Cang & Feldheim, 2013). Such a continuous organization preserves spatial relationships of the visual world. Contrary to retinotopy, the olfactory glomerular map consists of discrete units called glomeruli in which input neurons connect with the cognate output neurons based on neuronal type rather than soma position (Mombaerts et al., 1996; Gao et al., 2000; Vosshall et a, 2000). This discrete map represents a given odor by the combinatorial activation of specific glomeruli. Whereas continuous maps are readily built using gradients of guidance cues (Cang & Feldheim, 2013), how glomeruli are placed at specific locations in discrete maps is less clear (Murthy, 2011). Understanding the developmental origins of these neural maps is fundamental for deciphering the logic of their functional organization through which information is properly represented and processed.

The adult *Drosophila* olfactory map in the antennal lobe (equivalent of the vertebrate olfactory bulb) has proven to be a powerful model for studying mechanisms of wiring specificity, thanks to the type-specific connections between the presynaptic olfactory receptor neurons (ORNs) and the cognate postsynaptic projection neurons (PNs). Molecules and mechanisms first identified in this circuit have been found to play similar roles in the wiring of the mammalian brain (e.g., Hong et al., 2012; Berns et al., 2018; Pederick et al., 2021).

Assembly of the fly olfactory map begins with dendritic growth and patterning of PNs derived primarily from the anterodorsal (adPNs) and lateral (lPNs) lineages and born with an invariant birth order within each lineage (Jefferis et al., 2001, 2004; Marin et al., 2005; Yu et al., 2010; Lin et al., 2012) (**Figures 1A and B**). This patterning creates a prototypic olfactory map, prior to ORN axon innervation, indicative of the PN-autonomous ability to target dendrites into specific regions. However, earlier studies could only unambiguously follow the development of one single PN type – DL1 PNs (Jefferis et al., 2004). It remains unclear to date how the prototypic olfactory map is organized and what cellular mechanisms PN dendrites use to achieve targeting specificity (**Figure 1C_1–2_**). The initial map formation is further complicated by circuit remodeling during which embryonic-born PNs used in both the larval and adult circuits need to reorganize their neurites (Marin et al., 2005). How embryonic-born PNs coordinate remodeling with re-integration into the adult circuit is not known (**Figure 1C_3_**).

**Figure 1.**
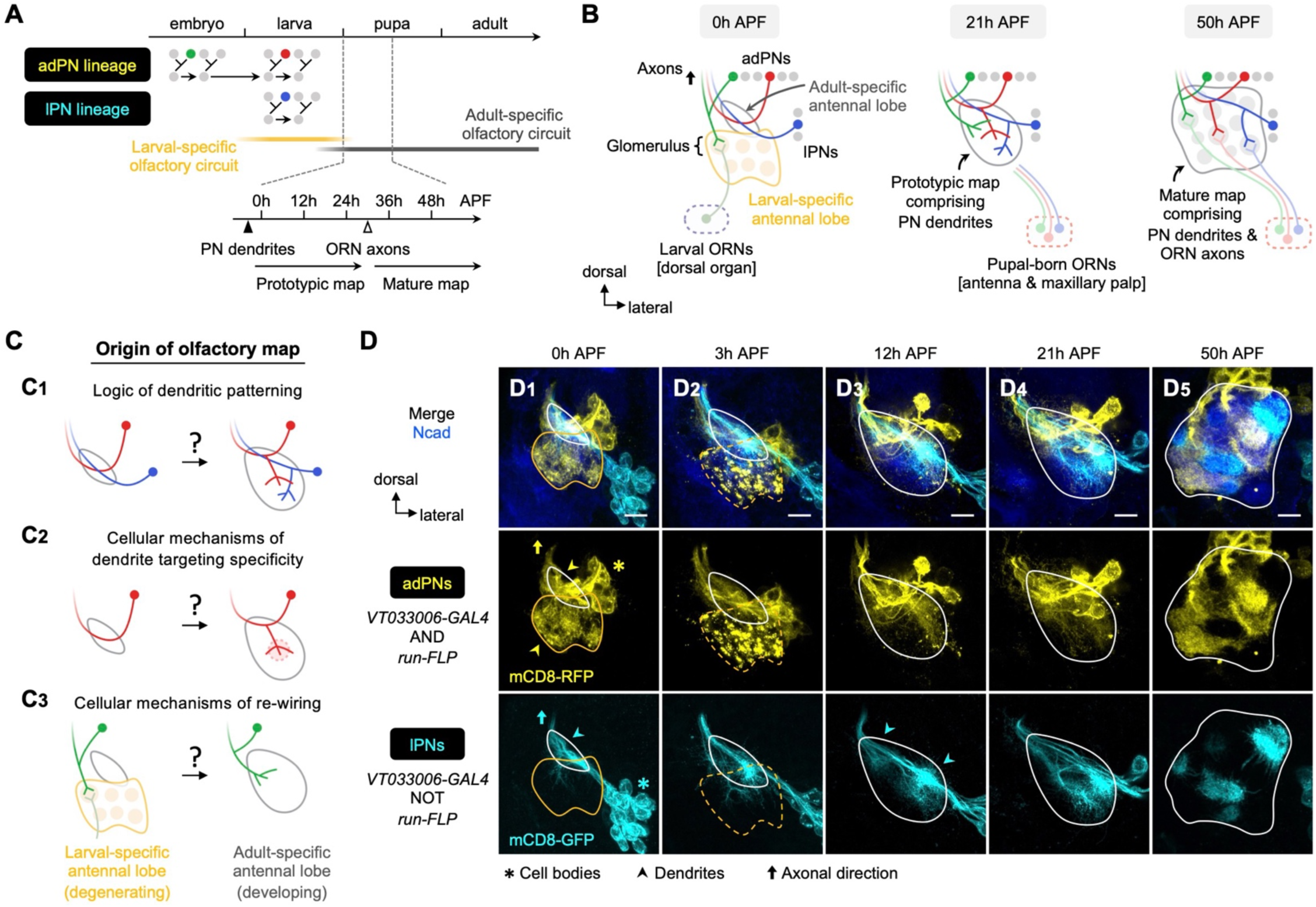
Organization and development of the adult olfactory circuit in *Drosophila*. **(A, B)** Timeline (**A**) and schematic illustration (**B**) of *Drosophila* olfactory circuit development. Green, red, and blue circles denote the birth of embryonic-born adPN, larval-born adPN, and larval-born lPN, respectively. At the onset of metamorphosis, larval-specific olfactory circuit degenerates; larval ORNs die while embryonic- born adPNs prune their larval-specific processes and re-extend new processes into the adult-specific olfactory circuit. In the adult-specific olfactory circuit, PN dendrites extend first and form a prototypic map. This is followed by extension of ORN axons and synaptic partner matching between cognate PN dendrites and ORN axons to form a mature map. Solid and open arrowheads in **A** indicate innervation timing of PN dendrites and ORN axons, respectively. **(C)** Overview of this study investigating the logic of dendritic patterning (**C_1_**; see Figures 3 and 4) as well as cellular mechanisms of dendrite targeting specificity (**C_2_**; see Figures 6 and 7) and re-wiring (**C_3_**; see Figure 8) that contribute to the developmental origin of the adult *Drosophila* olfactory map. **(D)** Staining of fixed brains at indicated stages showing dendrite development of adPNs (*VT033006+ run+*; labeled in yellow) and lPNs (*VT033006+ run–*; labeled in cyan). As *run-FLP* is expressed before 0h APF in adPN but not lPN neuroblasts, we can use it to label adPNs and lPNs with two distinct colors using an intersectional reporter (see **Materials and Methods** for the genotype). Yellow arrowheads in **D_1_** mark larval- and adult-specific dendrites of adPNs in larval- and adult-specific antennal lobes, respectively. Cyan arrowheads in **D_3_** denote specific targeting of lPN dendrites at the opposite ends of the dorsomedial- ventrolateral axis. **Common notations in this study:** Unless otherwise indicated, all images in this and subsequent figures are partial *z* projections of confocal stacks. Antennal lobe neuropils are revealed by N-Cadherin (Ncad; in blue) staining. Adult-specific (developing) antennal lobe is outlined with white solid line. Larval-specific antennal lobe is outlined with orange line (dashed line used to denote the degeneration stage) and is distinguished from the developing antennal lobe by the more intense nc82 staining as shown in **Figure 1 – figure supplement 1** (nc82 channel not shown here). Asterisks (*) indicate PN cell bodies, which are outside the antennal lobe neuropil (and sometimes appear on top because of the z-projections). Arrowheads mark PN dendrites. Arrows mark PN axons projecting towards higher olfactory centers (see **Figure 1 – figure supplement 2** for PN axons at their targets in the mushroom body and lateral horn). h APF: hours after puparium formation; h ALH: hours after larval hatching. DL: dorsolateral; DM: dorsomedial; VM: ventromedial; VL: ventrolateral. Scale bar = 10 µm.

Here, we set out to explore the origin of the olfactory map by performing a systematic and comparative study of PN dendrite development at type–specific resolution *in vivo*, and two- photon– and adaptive optical lattice light-sheet microscope–based time-lapse imaging of PN dendrites in early pupal brain explants. Our study uncovers wiring logic that directs PN dendrites to create an organized olfactory map, dendritic branch dynamics that lead to directional selectivity, and a novel re-wiring mechanism that facilitates timely olfactory map formation.

These wiring strategies used in the initial map organization lay the foundation of precise synaptic connectivity between PNs and ORNs in the final glomerular map.

## RESULTS

### Overview of *Drosophila* olfactory circuit development at lineage-specific resolution

We first described the development of *Drosophila* olfactory circuit using pupal brains double-labeled for adPNs and lPNs (**Figure 1D**; see genetic design in **Figure 2**). At the onset of metamorphosis (0 hours after puparium formation; 0h APF), the adult-specific antennal lobe (also referred to as ‘developing antennal lobe’) remained relatively small, located dorsolateral and posterior to the larval-specific antennal lobe (also referred to as ‘degenerating antennal lobe’) (**Figure 1D_1_**). As PN dendrites continued to grow and innervate the developing antennal lobe, its size increased considerably (**Figure 1D_1–3_**). By 12h APF, PNs already appeared to be sorting their dendrites into specific regions to form a prototypic map, as revealed by the heterogeneous patterning of lPN dendrites (arrowheads in **Figure 1D_3_**). From 21h to 50h APF, dendrites of adPNs and lPNs gradually segregated and eventually formed intercalated but non-overlapping glomeruli (**Figure 1D_4–5_**). The development of the adult-specific antennal lobe partially overlapped with the degeneration of the larval-specific antennal lobe, as indicated by fragmentation of the larval-specific dendrites of embryonic-born PNs at 3h APF (**Figure 1D_2_**). This gross characterization at the resolution of two PN lineages was consistent with earlier studies (Jefferis et al., 2004; Marin et al., 2005). However, the resolution was not sufficiently high to answer the questions we raised in Introduction (**Figure 1C**).

**Figure 2.**
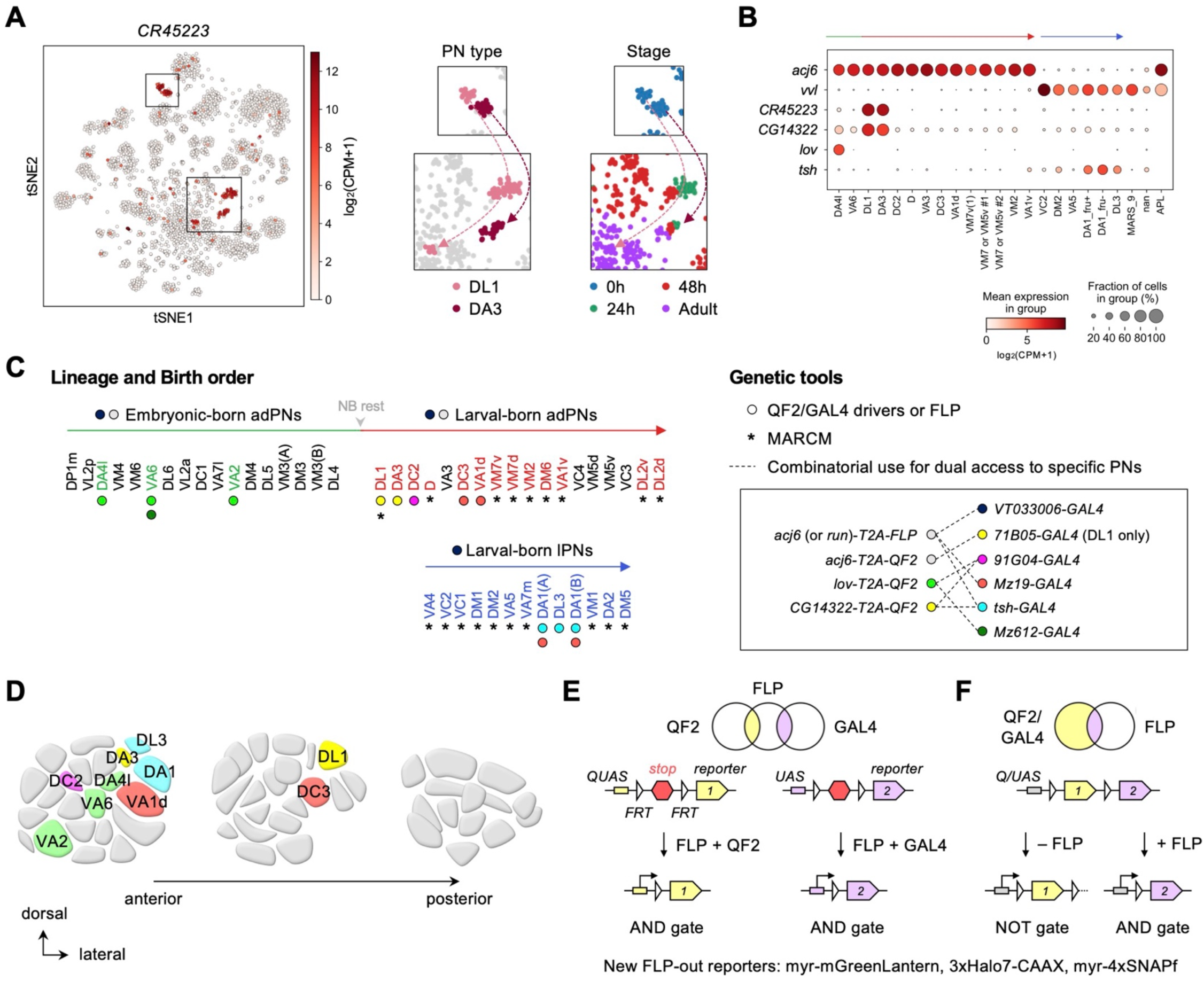
Expanded genetic toolkit for dual-color, type-specific labeling of PNs. **(A)** tSNE plot of PN single-cell transcriptomes, color-coded according to *CR45223* expression level in [log_2_(CPM+1)], where CPM stands for transcript counts per million reads. Zoom-in of boxes in the tSNE plot (left) are shown on the right, color-coded according to PN types and developmental stages. **(B)** Dot plot showing the expression of *acj6*, *vvl*, *CR45223*, *CG14322*, *lov*, and *tsh* in 0h APF PNs arranged according to their birth order and lineage (green: embryonic-born adPNs; red: larval-born adPNs; blue: larval- born lPNs). Unit of expression is [log_2_(CPM+1)] as in (A). **(C)** Birth orders of adPNs and lPNs [summarized from (Lin et al., 2012; Yu et al., 2010)] and genetic tools used to access them. **Left:** Accessible PN types are colored. Circles beneath the PN types denote *QF2/GAL4* drivers used to access them. Asterisks beneath the PN types denote access by MARCM. Grey arrowhead marks neuroblast (NB) rest. **Right:** Genetic tools. Inset shows combinatorial use of *QF2/FLP* and *GAL4* (linked by dashed lines) for comparative analyses of dendrite development of two groups of PNs in the same animal. **(D)** Schematic of glomerular projections of *QF2/GAL4-*accessible PNs in the adult antennal lobe. Indicated glomeruli are color-coded based on the genetic tools used to access them. See the color code in Figure 2C. **(E, F)** Schematic of intersectional logic gates for dual-color labeling of PNs. See **Figure 2 – figure supplement 2** for newly generated FLP-out reporters.

### Expanded genetic toolkit for type-specific labeling of PNs during early pupal development

To reveal how PN dendrites initiate olfactory map formation at high spatiotemporal resolution, we needed genetic access to specific PN types during early pupal development. From our recently deciphered single-cell PN transcriptomes (Xie et al., 2021), we searched for genetic markers that are expressed strongly and persistently in single or a few PN types across pupal development. This transcriptome-instructed search led to identification of *CR45223* (in place of this non-coding gene, we used the adjacent *CG14322* that exhibits nearly identical expression pattern), *lov*, and *tsh* (**Figures 2A and B**; **Figure 2 – figure supplement 1**).

Next, using CRISPR/Cas9, we generated knock-in transgenic QF2 expression driver lines in which *T2A-QF2* (or *T2A-FLP* for intersection) was inserted immediately before the stop codon of the endogenous gene (**Figure 2 – figure supplement 2**). The self-cleaving peptide T2A allows QF2 to be expressed in the same pattern as the endogenous gene (Diao & White, 2012).

With these new *QF2* lines together with existing *GAL4* lines that label additional PN types (Xie et al., 2019), we now have an expanded toolkit accessing PNs ranging from early- to late-born PNs, from adPN to lPN lineages, and from PNs with neighboring glomerular projections to those with distant projections in the adult antennal lobe (**Figures 2C and D**). As QF2*/QUAS* and GAL4*/UAS* expression systems operate orthogonally to each other (Potter et al., 2010), we crossed our *QF2* lines with existing *GAL4* lines for simultaneous labeling of distinct PN types in the same brain (see inset in **Figure 2C**). This combinatorial use of driver lines permitted comparative analyses of development of distinct PN types with minimal biological and technical variations.

To limit driver expression only in PNs, we applied intersectional logic gates (AND and NOT gates) using our newly generated conditional reporters genetically encoding either mGreenLantern, Halo tags, and/or SNAP tags (Kohl et al., 2014; Sutcliffe et al., 2017; Campbell et al., 2020) (**Figures 2E and F; Figure 2 – figure supplement 3**). These reporters can be broadly used in other systems. Finally, we used MARCM (Lee & Luo, 1999) to label PNs that remain inaccessible due to lack of drivers (**Figure 2C**; discussed in **Figure 3**).

**Figure 3.**
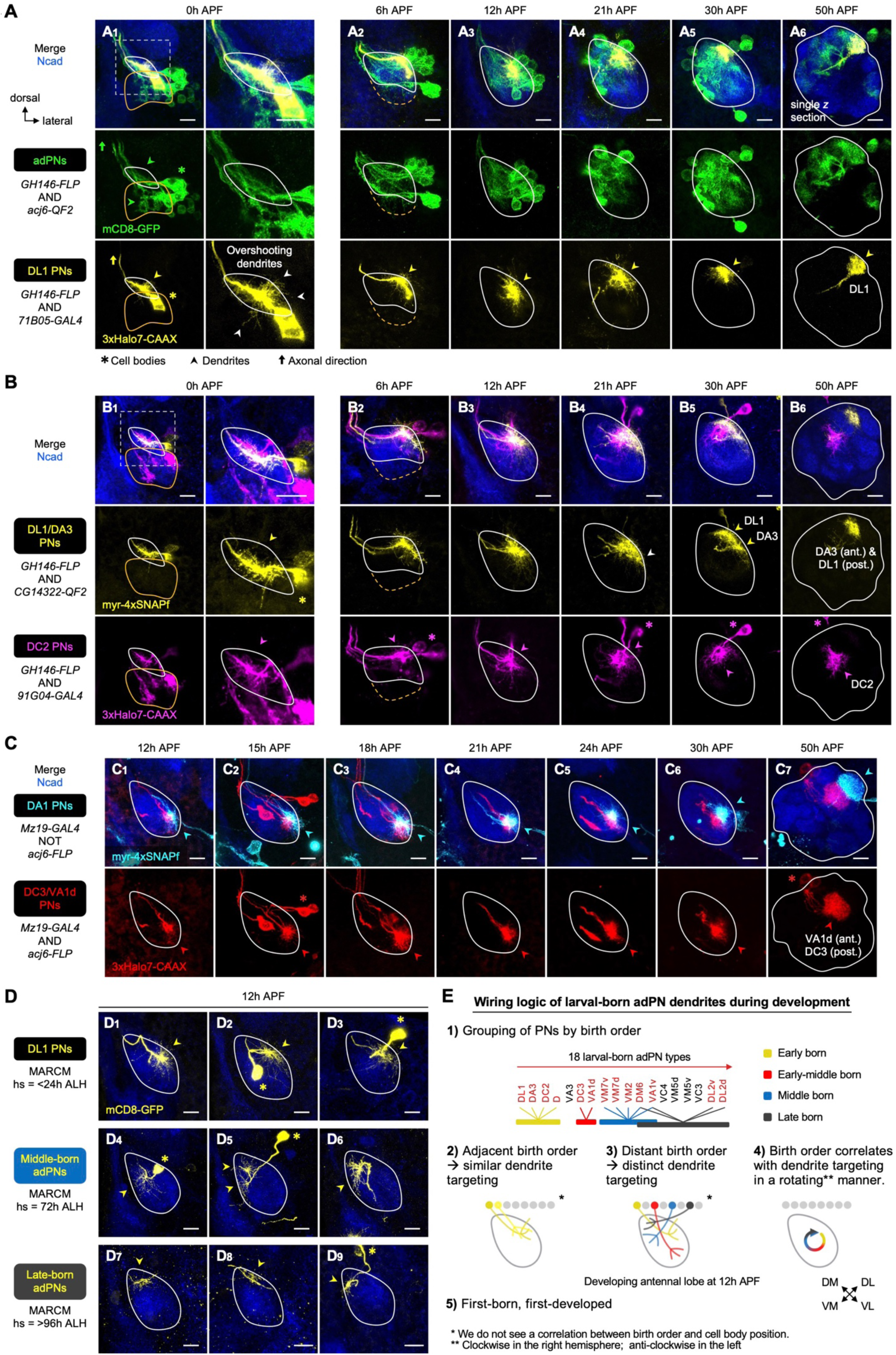
Birth order–dependent spatial patterning of adPN dendrites in the developing antennal lobe. **(A)** Confocal images of fixed brains at indicated stages showing dendrite development of adPNs (*acj6+*; labeled in green) and DL1 adPNs (*71B05+*; labeled in yellow). Right column of A_1_ shows zoom-in of the dashed box. The labeling of *acj6+* adPNs outlines the developing antennal lobe and is used in dual-color AO-LLSM imaging later (see Figure 7A**–C**). White arrowheads in **A**_1_ mark dendrites overshooting the antennal lobe. **(B)** Confocal images of fixed brains at indicated stages showing dendrite development of DL1/DA3 adPNs (*CG14322+*; labeled in yellow) and DC2 adPNs (*91G04+*; labeled in magenta). As *91G04-GAL4* labels some embryonic-born PNs from 0–6h APF, their neurites are found in the larval-specific antennal lobe (**B_1, 2_**). Right column of **B_1_** shows zoom-in of the dashed box. White arrowhead in **B_4_** denotes the more ventrally targeted DL1/DA3 dendrites. **(C)** Confocal images of fixed brains at indicated stages showing dendrite development of DC3/VA1d adPNs (*Mz19+ acj6+*; labeled in red) and DA1 lPNs (*Mz19+ acj6–*; labeled in cyan). **(D)** Confocal images of single-cell MARCM clones (in yellow) of DL1 PNs (**D_1–3_**), middle larval-born adPNs (**D_4–6_**) and late larval-born adPNs (**D_7–9_**) in 12h APF pupal brains, generated by heat shocks (hs) at indicated times. **(E)** Summary of wiring logic of larval-born adPN dendrites to form olfactory map in the 12h APF developing antennal lobe. See Figure 1 legend for common notations.

### Early larval-born adPN dendrites initially share similar targeting regions

Using the new genetic tools, we first re-visited the dendrite development of DL1 PNs—the first larval-born adPN type—using pupal brains double-labeled for DL1 PNs (labeled by *71B05- GAL4*) and adPNs (**Figure 3A**). Consistent with our previous study (Jefferis et al., 2004), DL1 PNs already showed robust dendritic growth at the wandering third instar larval stage (**Figure 3 – figure supplement 1A**). At 0h APF, DL1 PN dendrites extended radially outwards from the main process, reaching nearly the entire developing antennal lobe and often overshooting it (white arrowheads in **Figure 3A_1_**), likely surveying the surroundings. By 6h APF, most of the dendrites already occupied the dorsolateral (DL) corner of the antennal lobe (**Figure 3A_2_**). As the antennal lobe continued to grow, this dorsolateral positioning of the DL1 PN dendrites remained largely unchanged (**Figure 3A_3–6_**). From 21h APF onwards, the dendrites underwent progressive refinement: they were restricted into a smaller area by 30h APF (**Figure 3A_4–5_**), and eventually formed a compact, posterior glomerulus by 50h APF (**Figure 3A_6_** showing a single *z* section).

To assess whether other PN types follow the same developmental trajectory, we next examined *CG14322*+ PNs, which include DL1 PNs and DA3 PNs—the first and second larval-born adPN types, respectively. In the same brain, we also labeled with a different fluorophore DC2 PNs—the third larval-born adPN type (**Figure 3B**). The dendritic pattern of DL1/DA3 PNs appeared indistinguishable from that of DL1 PNs from 0h to 12h APF (compare yellow channel of **Figure 3B_1–3_** with **Figure 3A_1–3_**), suggesting that DL1 and DA3 PN sent dendrites to the same region in the antennal lobe. We began to see differences in 21h APF pupal brains in which DL1/DA3 PN dendrites not only occupied the dorsolateral region but also spread ventrally (white arrowhead in **Figure 3B_4_**; compare with **Figure 3A_4_**). The more ventrally targeted dendrites likely belong to DA3 PNs. This suggests that ∼21h APF marks the beginning of dendritic segregation of DL1 and DA3 PNs. By 30h APF, DL1 and DA3 dendrites were clearly separable (**Figure 3B_5_**), which respectively formed more posteriorly and anteriorly targeted glomeruli at 50h APF (**Figure 3B_6_**; see single *z* sections in **Figure 3 – figure supplement 1C**).

Next, we focused on the third-born—DC2 PNs labeled by *91G04-GAL4* (**Figure 3B**). This *GAL4* labeled additional embryonic-born adPNs from 0h to 6h APF, but the expression in these PNs diminished afterwards. As embryonic-born adPNs do not have any dendrites in the developing antennal lobe at 0h APF (discussed in **Figure 7**), dendrites found in the antennal lobe should belong to the larval-born DC2 PNs. Like DL1/DA3 PNs, DC2 PNs initiated radial dendritic extension across the antennal lobe at 0h APF (**Figure 3B_1_; Figure 3 – figure supplement 1B**). Notably, DL1/DA3 and DC2 PN dendrites exhibited substantial overlap from 0h to 12h APF and shared a similar targeting region at the dorsolateral corner from 6h to 12h APF (**Figure 3B_1–3_**). It was not until 21h APF that DL1, DA3, and DC2 dendrites began to segregate from each other along both medial-lateral and anterior-posterior axes (**Figure 3B_4–5_**). By 50h APF, DC2 glomerulus was separated from DL1/DA3 glomeruli by intermediate glomeruli (**Figure 3B_6_**).

In summary, dendrites of consecutively larval-born DL1, DA3, and DC2 adPNs (here collectively named ‘early larval-born adPNs’) develop in a similar fashion and share a similar targeting region at early pupal stages (0–12h APF). This is then followed by their segregation into distinct regions close to their adult glomerular positions during mid-pupal stages (21–50h APF).

### Larval-born adPNs with distant birth order send dendrites to distinct regions

The analysis of early larval-born adPNs (**Figures 3A and B**) led us to hypothesize that larval-born adPNs might use their birth order to coordinate dendrite targeting during early pupal stages. If this were true, we would expect dendrites of larval-born adPNs with distant birth order to occupy distinct regions. To test this hypothesis, we compared dendrite targeting regions of early larval-born adPNs with those of later-born adPNs.

We first examined DC3/VA1d adPNs (referred to as ‘early-middle larval-born adPNs’) using *Mz19-GAL4* (**Figure 3C**). This *GAL4* is expressed in 3 PN types from 24h APF to adulthood: DC3 adPNs, VA1d adPNs, and DA1 lPNs (Jefferis et al., 2004). To distinguish adPNs from lPNs, we previously adopted a FLP-out strategy labeling *Mz19+* PNs with either GFP or RFP based on their lineages and studied dendrite segregation and refinement during mid-pupal stages (Li et al., 2021) (**Figure 3C_4–7_**). However, the weak *GAL4* expression before 24h APF prevented us from visualizing any dendrites at earlier stages. To overcome this, we incorporated Halo and SNAP chemical labeling (Kohl et al., 2014) in place of the immunofluorescence approach. This modification substantially extended the detection to developmental stages as early as 12h APF (**Figure 3C_1_**). We found that, from 12h to 21h APF, DC3/VA1d PN dendrites targeted the ventrolateral (VL) corner of the antennal lobe (**Figure 3C_1–4_**). Thus, early (DL1/DA3/DC2) and early-middle (DC3/VA1d) larval-born adPN dendrites occupy distinct regions at 12h APF.

As we did not have reliable drivers to access other later-born PNs at early pupal stages, we turned to MARCM (Lee & Luo, 1999) to generate heat shock-induced single-cell clones of PNs born at different times (**Figure 3 – figure supplement 2**). We used *GH146-GAL4(IV)*, a PN driver that labels the majority of PN types, including later-born adPNs (**Figure 3 – figure supplement 2D–E**), with a tight temporal control of heat shock and analyzed heat shock-induced animals that were among the first to form puparium to minimize the effects of unsynchronized development among individual animals (see **Materials and Methods** for details). These optimizations permitted a systematic clonal analysis at higher PN type-specific resolution that correlates with birth time. Applying heat shocks at specific times, we accessed four groups of larval-born adPNs: (1) first-born (DL1), (2) early-born (DL1, DA3, DC2 and D), (3) middle-born (VM7v, VM7d, VM2, DM6 and VA1v), and (4) late-born (DM6, VA1v, DL2v, DL2d). We note that DM6 and VA1v PNs belonged to both groups of middle- and late-born adPNs, reflecting the nature of short birth timing differences among these later-born PNs. Using this strategy, we could also label lPNs born at different times (**Figure 3 – figure supplement 2F**).

Clonal analysis revealed that, at 12h APF, the first-born DL1 adPNs sent dendrites to the dorsolateral corner of the antennal lobe as expected (**Figure 3D_1–3_**). By contrast, dendrites of middle larval-born adPNs occupied a large region on the medial/dorsomedial (M/DM) side (**Figure 3D_4–6_**). The dendritic arborization patterns of these PNs varied widely, most likely because they belonged to different PN types. Intriguingly, late larval-born adPN dendrites targeted the <u>p</u>eripheral, <u>d</u>orso<u>m</u>edial (abbreviated as pDM) corner where the staining of the pan- neuropil marker N-Cadherin was relatively weak (**Figure 3D_7–9_**). The weak staining on the periphery implies that this area is less populated by PN dendrites (the major constituent of the antennal lobe neuropil at this stage), possibly because (1) this area is not innervated by many PNs and/or (2) the dendrites of late-born PNs innervate later and remain less elaborate than earlier-born PNs (we will explore this later).

Together, our data (**Figure 3A–D**) suggest that larval-born adPNs with adjacent birth order send dendrites to similar regions of the developing antennal lobe whereas those with distant birth order send dendrites to distinct regions (**Figure 3E_2, 3_**). We therefore propose that, at early pupal stages, larval-born adPNs are grouped by their birth order, and those belonging to the same group share similar dendrite targeting specificity (**Figure 3E_1_**). Notably, the birth order of the examined PNs does not specify dendrite targeting randomly (**Figure 3E_4_**). Rather, the stereotyped dendritic pattern in the prototypic map correlates with the birth order in an organized manner (rotating clockwise in the right hemisphere when viewed from the front; anti-clockwise in the left: early↔ DL; early-middle↔VL; middle↔M/DM; late↔pDM). One can therefore infer at least the approximate birth order of a larval-born adPN based on its initial dendrite targeting, and *vice versa*.

### Timing of larval-born adPN dendrite targeting depends on birth order

Having provided evidence for birth order–dependent spatial patterning of larval-born adPN dendrites, we next asked whether the timing of dendritic extension and targeting is also influenced by birth order. We noticed that the extent of dendritic innervation of 0h APF first-born DL1 adPNs resembled that of 6h APF middle-born adPNs (compare **Figure 3 – figure supplement 3A_1–4_** with **Figure 3 – figure supplement 3B_5–8_**). Such a resemblance was also seen between 0h APF middle-born and 6h APF late-born adPNs (compare **Figure 3 – figure supplement 3B_1–4_** with **Figure 3 – figure supplement 3C**). Quantitative analyses of the exploring volume of dendrites and the number of terminal branches showed that, at 0h APF, DL1 PN dendrites were more elaborate than middle-born PN dendrites (**Figure 3 – figure supplement 3F**). By 6h APF, the middle-born appeared to catch up, showing an extent of innervation comparable to DL1 PNs.

We next examined when the dendrites reach their targeting regions. We found that whereas early larval-born adPNs (DL1, DA3, DC2) concentrated their dendrites to the dorsolateral corner by 6h APF (**Figure 3B_2_; Figure 3 – figure supplement 3A_5–8_**), later-born PNs concentrated their dendrites to the medial/dorsomedial or peripheral dorsomedial side at 12h APF (**Figure 3D_4–9_; Figure 3 – figure supplement 3B_5–8_–C**). Thus, our results suggest larval-born adPN dendrites innervate and pattern the antennal lobe using a ‘first born, first developed’ strategy.

### Contribution of lineage to early PN dendritic patterning

Both lineage and birth order of PNs contribute to the eventual glomerular choice of their dendrites (Jefferis et al., 2001). What is the involvement of lineage in the prototypic map formation? Do lPN dendrites pattern the developing antennal lobe following similar rules as adPNs? To characterize lPN dendrite development at type–specific resolution, we used *tsh-GAL4* to genetically access DA1/DL3 lPNs, and MARCM clones of lPNs as a complementary approach (**Figure 4**). We focused on the dendritic patterns of *tsh+* DA1/DL3 lPNs from 0h to 12h APF as *tsh-GAL4* labeled additional PNs from 21h APF onwards (**Figure 4A_4–6_; Figure 4 – figure supplement 1B_4–6_; Figure 4 – figure supplement 2; Figure 2 – figure supplement 1**).

**Figure 4.**
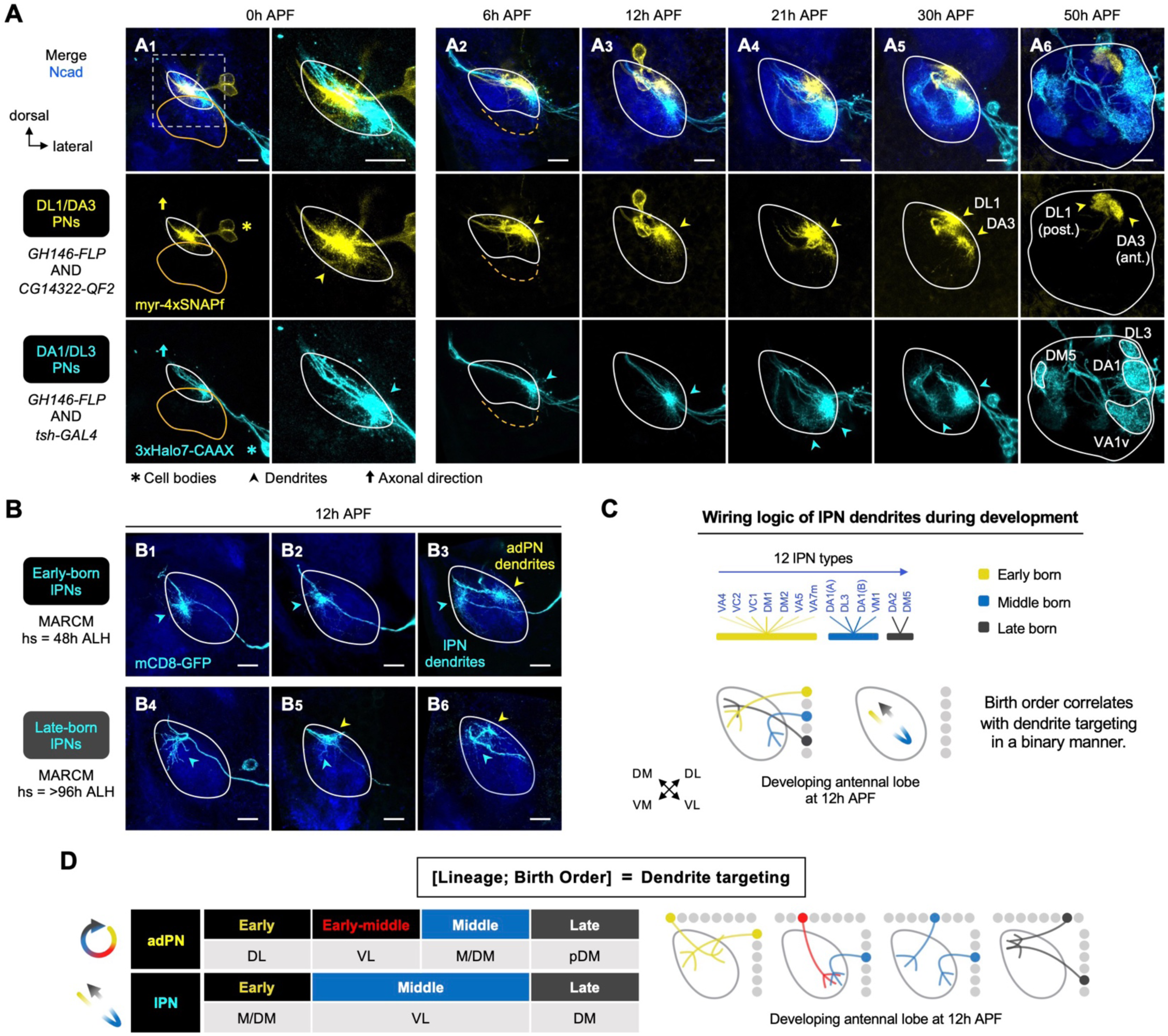
Birth order–dependent spatial patterning of lPN dendrites in the developing antennal lobe. **(A)** Confocal images of fixed brains at indicated stages showing dendrite development of DL1/DA3 adPNs (*CG14322+*; labeled in yellow) and DA1/DL3 lPNs (*tsh+*; labeled in cyan). Right column of A_1_ shows zoom-in of the dashed box. **(B)** MARCM clones (in cyan) of early (**B_1–3_**) and late (**B_4–6_**) larval-born lPNs in 12h APF pupal brains, generated by heat shocks (hs) at indicated times. In **B_3_**, **B_5_**, and **B_6_**, single-cell clones of adPN (yellow arrowheads) and lPN (cyan arrowheads) lineages were simultaneously labeled. **(C)** Summary of wiring logic of larval-born lPN dendrites to form olfactory map in the 12h APF developing antennal lobe. **(D)** Summary of determination of dendrite targeting of larval-born PNs by lineage and birth order. See Figure 1 legend for common notations.

Examination of pupal brains double-labeled with DA1/DL3 lPNs (referred to as ‘middle larval-born lPNs’) and DL1/DA3 adPNs revealed that, like the early larval-born adPNs, dendritic growth of DA1/DL3 lPNs was evident by the wandering third instar larval stage (**Figure 4 – figure supplement 1A)**. At this stage, most DA1/DL3 lPN dendrites innervated the antennal lobe and intermingled with those of DL1/DA3 adPNs. From 0h to 12h APF, despite a high degree of overlap among those dendrites that explored the surroundings, DA1/DL3 lPN dendrites primarily targeted an area ventrolateral to those of DL1/DA3 adPNs (**Figure 4A_1–3_**). Such a spatial distinction was also observed between middle larval-born adPNs and lPNs in 0h and 6h APF pupal brains where occasionally single-cell clones from both lineages were simultaneously generated by MARCM (**Figure 3 – figure supplement 3D_1–4, 7–10_**). Thus, at least some adPNs and lPNs sort their dendrites into distinct regions very early on regardless of birth timing.

Next, we used MARCM to ask if lPNs born earlier and later than DA1/DL3 lPNs would send dendrites to regions different from that of DA1/DL3 lPNs. We found that dendrites of early- born lPNs primarily occupied the medial/dorsomedial side of the antennal lobe (**Figure 4B_1–3_)**; we note that adPNs born at the same time sent dendrites to the dorsolateral side (see yellow arrowhead in **Figure 4B_3_)**. Also, in contrast to the ventrolateral targeting of middle-born lPN dendrites, late-born lPNs sent dendrites to the dorsomedial corner (**Figure 4B_4–6_)**. Like larval-born adPNs, late-born lPNs innervated the antennal lobe later than earlier-born lPNs (**Figure 3 – figure supplement 3D_7–12_–E, G**).

These data suggest that, at early pupal stages, lPN dendrites pattern the developing antennal lobe following similar rules as larval-born adPNs: grouping by birth order; adjacent birth order → similar dendrite targeting; distant birth order → distinct dendrite targeting; ‘first born, first developed’. However, unlike the correlation of birth order and target positions in a rotational manner for adPNs (**Figure 3E**), the lPN dendritic map formation appears binary: early↔M/DM; middle↔VL; late↔DM (**Figure 4C**). Our type-specific characterization corroborated with the gross examination of the lPN dendrites as previously reported (Jefferis et al., 2004): at 12h APF, lPN dendrites mostly occupied the opposite corners along the dorsomedial-ventrolateral axis, leaving the middle of the axis largely devoid of lPN dendrites (arrowheads in **Figure 1D_3_**).

In summary, lineage and birth order of larval-born PNs contribute to their dendrite targeting in a combinatorial fashion (**Figure 4D**). The wiring logic of PN dendrites in the developing antennal lobe can therefore be represented by [lineage, birth order] = dendrite targeting; one can deduce the unknown if the other two are known.

### An explant system for time-lapse imaging of PN development at early pupal stages

So far, we have identified wiring logic governing the initial dendritic map formation (**Figures 3 and 4**). To examine dendrite targeting at higher spatiotemporal resolution, we established an early-pupal brain explant culture system based on previous protocols (Özel et al., 2015; Rabinovich et al., 2015; Li and Luo, 2021; Li et al., 2021), and performed single- or dual-color time-lapse imaging with two-photon microscopy as well as adaptive optical lattice light-sheet microscopy (AO-LLSM) (**Figure 5A–C**). The following lines of evidence support that our explant system recapitulates key features of *in vivo* olfactory circuit development.

**Figure 5.**
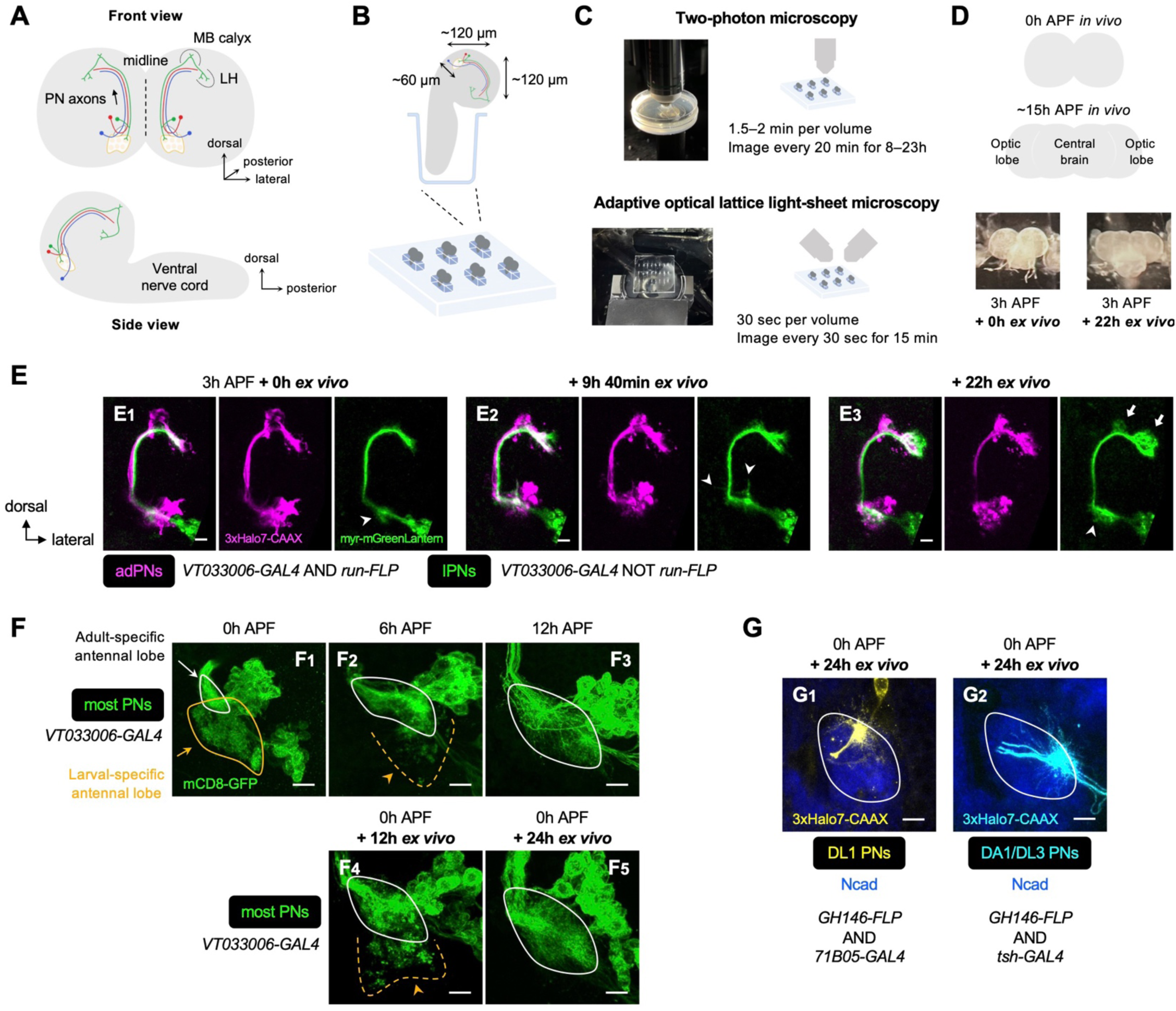
Establishment of explant system for time-lapse imaging of olfactory map formation. **(A)** Schematic of anatomical organization of the olfactory circuit in early pupal brain (0–3h APF). Green, red, and blue denote embryonic-born adPN, larval-born adPN and larval-born lPN, respectively. MB: mushroom body; LH: lateral horn. **(B)** Schematic of explant culture system for early pupal brains. Wells created in the Sylgard plate from which brains were imbedded are shown in blue. **(C)** Schematic of explant culture and imaging system for early pupal brains. **(D) Top:** Schematic of morphological changes of brain lobes from 0h to ∼15h APF during normal development. **Bottom:** Morphologies of a brain explant dissected at 3h APF and cultured for 0h *ex vivo* and that cultured for 22h *ex vivo*. **(E)** Two-photon time-lapse imaging of adPNs (*VT033006+ run*+; labeled in magenta) and lPNs (*VT033006+ run*–; labeled in green) in pupal brain dissected at 3h APF and cultured for 0–22h *ex vivo.* Arrowheads mark dynamic but transient dendritic protrusions of lPNs in **E_1, 2_**, and extensive dendritic innervation of lPNs in **E_3_**. Arrows in **E_3_** mark axonal innervation of lPNs in mushroom body calyx and lateral horn. **(F)** Confocal images of antennal lobes labeled by *VT033006+* PNs (in green) at 0h (**F_1_**), 6h (**F_2_**) and 12h (**F_3_**) APF *in vivo*. Confocal images of antennal lobes labeled by *VT033006+* PNs in pupal brains dissected at 0h APF and cultured for 12h (**F_4_**) and 24h (**F_5_**) *ex vivo*. **(G)** Dendrite targeting regions of DL1 PNs (*71B05+*; in yellow; **G_1_**) and DA1/DL3 PNs (*tsh+*; in cyan; **G_2_**) in the antennal lobes in pupal brains dissected at 0h APF and cultured for 24h *ex vivo.* Antennal lobes are revealed by N-Cadherin (Ncad; in blue) staining. See Figure 1 legend for common notations.

First, during normal development, the morphology of the brain lobes changes from spherical at 0h APF to more elongated rectangular shapes at 15h APF (Rabinovich et al., 2015). After 22h *ex vivo* culture, the spherical hemispheres of brains dissected at 3h APF became more elongated, mimicking ∼15h APF *in vivo* brains characterized by the separation of the optic lobes from the central brain (**Figure 5D**).

Second, dual-color, two-photon imaging of PNs every 20 min for 22h revealed that lPNs in 3h APF brains initially produced dynamic but transient dendritic protrusions in many directions, followed by extensive innervation into the antennal lobe (arrowheads in **Figure 5E_1–3_**; **Figure 5 – video 1**). In higher brain centers, lPN axons clearly showed direction-specific outgrowth of collateral branches into the mushroom body calyx as well as forward extension into lateral horn (arrows in **Figure 5E_3_**), thus resembling *in vivo* development (**Figure 1 – figure supplement 2**).

Third, larval-specific dendrites observed in 0h APF brains cultured for 12h *ex vivo* (orange arrowhead in **Figure 5F_4_**) were no longer seen in those cultured for 24h *ex vivo* (**Figure 5F_5_**), indicative of successful pruning and clearance of larval-specific dendrites. Also, the size of the developing antennal lobe in the brains cultured for 24h *ex vivo* increased considerably (**Figure 5F_5_**). These imply that olfactory circuit remodeling (degeneration of larval-specific processes and growth of adult-specific processes) proceeds normally, albeit at a slower rate (compare with **Figure 5F_1–3_**).

Fourth, dendrites from genetically identified DL1 and DA1/DL3 PNs targeted to their stereotyped locations in the antennal lobe in 0h APF brains cultured for 24h *ex vivo*, (**Figure 5G**), mimicking *in vivo* development (**Figure 4A**).

Finally, segregation of dendrites of PNs targeting to neighboring proto-glomeruli could be recapitulated in brains dissected at 24h APF and cultured for 8h (**Figure 5 – figure supplement 1**; **Figure 5 – video 2**). Specifically, despite constant dynamic interactions among dendrites that explore the surroundings (arrowheads in **Figure 5 – figure supplement 1A_2–4_**), DC3/VA1d and DA1 PNs exhibited a 1–2 µm increase in the distance between centers of the two dendritic masses and a substantial decrease in the overlap of their core targeting regions (**Figure 5 – figure supplement 1B–D**). Taken together, these data support that the explant culture and imaging system established here reliably captures key neurodevelopmental events starting from early pupal stages.

### Single-cell, two-photon imaging reveals active dendrite targeting

Our observation in fixed brains revealed that dendrites of DL1 adPNs transition from a uniform extension in the antennal lobe at 0h APF to concentration at the dorsolateral corner of the antennal lobe at 6h APF (**Figure 3A**). To identify mechanisms of dendrite targeting specificity that could be missed in static developmental snapshots, we performed two-photon time-lapse imaging of single-cell MARCM clones of DL1 PNs in 3h APF brains (**Figure 6**; **Figure 6 – figure supplement 1**; **Figure 6 – video 1**). Although we did not have a counterstain outlining the antennal lobe, we could use the background signals to discern the orientation of DL1 PNs in the brain (**Figure 6 – figure supplement 1A**). The final targeting regions relative to the antennal lobe revealed by *post hoc* fixation and immunostaining confirmed proper dendrite targeting (yellow arrowhead in **Figure 6A_10_**; **Figure 6 – figure supplement 1B–C**).

**Figure 6.**
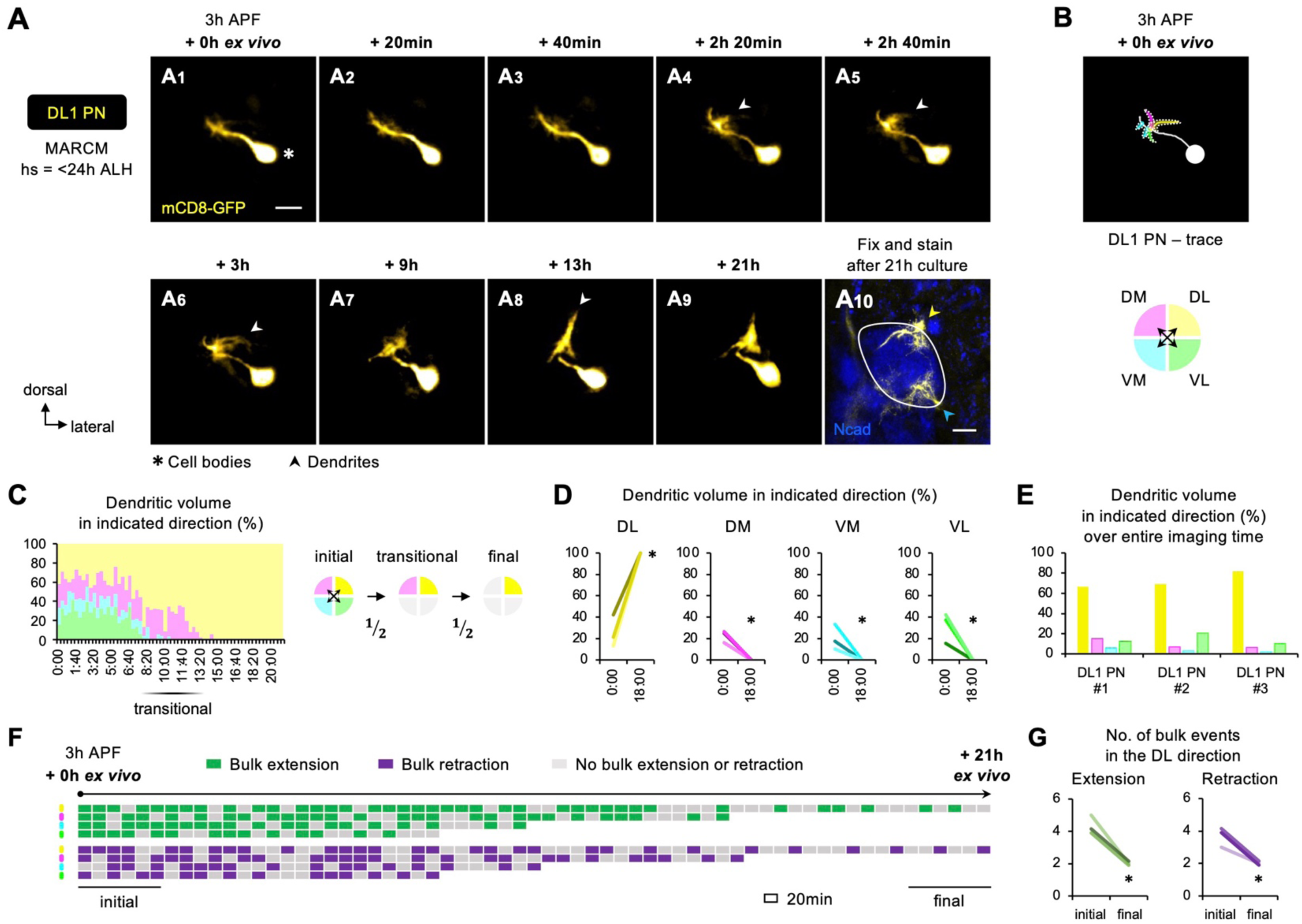
Two-photon time-lapse imaging reveals active dendrite targeting. **(A)** Two-photon time-lapse imaging of MARCM-labeled DL1 PN (pseudo-colored in yellow) in a brain dissected at 3h APF and cultured for 21h *ex vivo* (**A_1–9_)**. Arrowheads in **A_4–6_** denote protrusions of dendritic branches towards the dorsolateral direction. After 21h culture, explant was fixed and immune-stained for N- Cadherin (Ncad; in blue) to outline the developing antennal lobe (**A_10_)**. Yellow and cyan arrowheads indicate DL1 PN dendrites and processes of other *GH146+* cells, respectively. **(B)** Neurite tracing of DL1 PN at the beginning of live imaging (3h APF + 0h *ex vivo*). Dendrites are categorized based on the directions to which they extend and color-coded accordingly. **(C)** Left: Quantification of the percentage of dendritic volume in indicated direction during the time-lapse imaging period reveals a transitional phase during which dendrites were found in only two out of the four directions. Right: Schematic of the initial, transitional and final phases during the course of targeting. ‘½’ denotes the reduction of available trajectory directions by half. Timestamp 00:00 refers to HH:mm; H, hour; m, minute. **(D)** Quantification of the percentage of DL1 PN dendritic volume in indicated direction in 3h APF cultured brains at the beginning (0h *ex vivo*) and at/near the end of imaging (18h *ex vivo*). DL1 PN sample size = 3. *t* test; *\**, *p* < 0.05. Timestamp 00:00 refers to HH:mm; H, hour; m, minute. **(E)** Quantification of the percentage of sum of DL1 PN dendritic volume in indicated directions throughout the entire imaging time. DL1 PN sample size = 3. **(F)** Bulk dendrite dynamics of DL1 PN in Figure 6A. Each row represents bulk dendritic dynamics in the indicated direction (color coded as in Figure 6B) across the 22-h imaging period. Each block represents a 20- min window. Bulk extension (in green) and retraction (in magenta) events are defined as dendrites extending and retracting more than 2 μm between two consecutive time windows. The first and last 6 consecutive windows refer to the initial and final phases of imaging. **(G)** Quantification of number of bulk extension and retraction events in the dorsolateral direction during the initial and final phases of imaging. DL1 PN sample size = 3. *t* test; *\**, *p* < 0.05.

Using DL1 PN in **Figure 6A** (pseudo-colored in yellow; **Figure 6 – video 1**) as an example, we observed that the PN initially extended dendrites in every direction (**Figure 6A_1–3_**), like what we observed in fixed tissues (**Figure 3A_1_**). The first sign of active targeting emerged at 2h 20min *ex vivo* when DL1 PN began to generate long, albeit transient, dendritic protrusions in the dorsolateral direction; these selective protrusions were more prominent at 3h *ex vivo* (arrowheads in **Figure 6A_4–6_**). The dorsolateral targeting continued to intensify, leading to the formation a highly focal dendritic mass seen at 13h *ex vivo* (arrowhead in **Figure 6A_8_**). As the dendrites reached the dorsolateral corner and explored locally, the change in shape appeared less pronounced (**Figure 6A_9_**).

To quantitatively characterize the active targeting process, we categorized the bulk dendritic masses emanated from the main process according to their targeting directions: DL, DM, VM, and VL (**Figure 6B**). During the initial phase, the percentage of dendritic volume in each direction varied from 10% to 40% (**Figures 6C and D)**, indicative of active exploration with little targeting specificity. Despite these variations, the total amount of dendritic mass seen in the VM direction over the entire imaging time (area under graph of **Figure 6C**) was the smallest across all samples examined (**Figure 6E)**. The initial phase of exploration in every direction was followed by a ∼4-hour transitional phase during which DL1 PNs predominantly extended dendrites in 2 of the 4 directions (**Figure 6C**; **Figure 6 – figure supplement 1D–E**). One of the 2 directions was always DL whereas the other was either DM or VL but never VM. At the final phase, DL1 PN dendrites always preferred DL out of the 2 available directions.

Lastly, we analyzed the bulk dendritic movements. We defined bulk extension and retraction events when dendrites respectively extended and retracted more than 2 μm between two consecutive time frames. The analyses showed a striking shift from frequent extension and retraction towards stabilization, reflecting the pre- and post-targeting dynamics respectively (**Figure 6F and G)**.

Hence, long-term two-photon imaging of single-cell DL1 PNs revealed that dendrite targeting specificity increases over time via active targeting in specific direction and stepwise elimination of unfavorable trajectory choices (see summary in **Figure 7F_1–3_)**.

**Figure 7.**
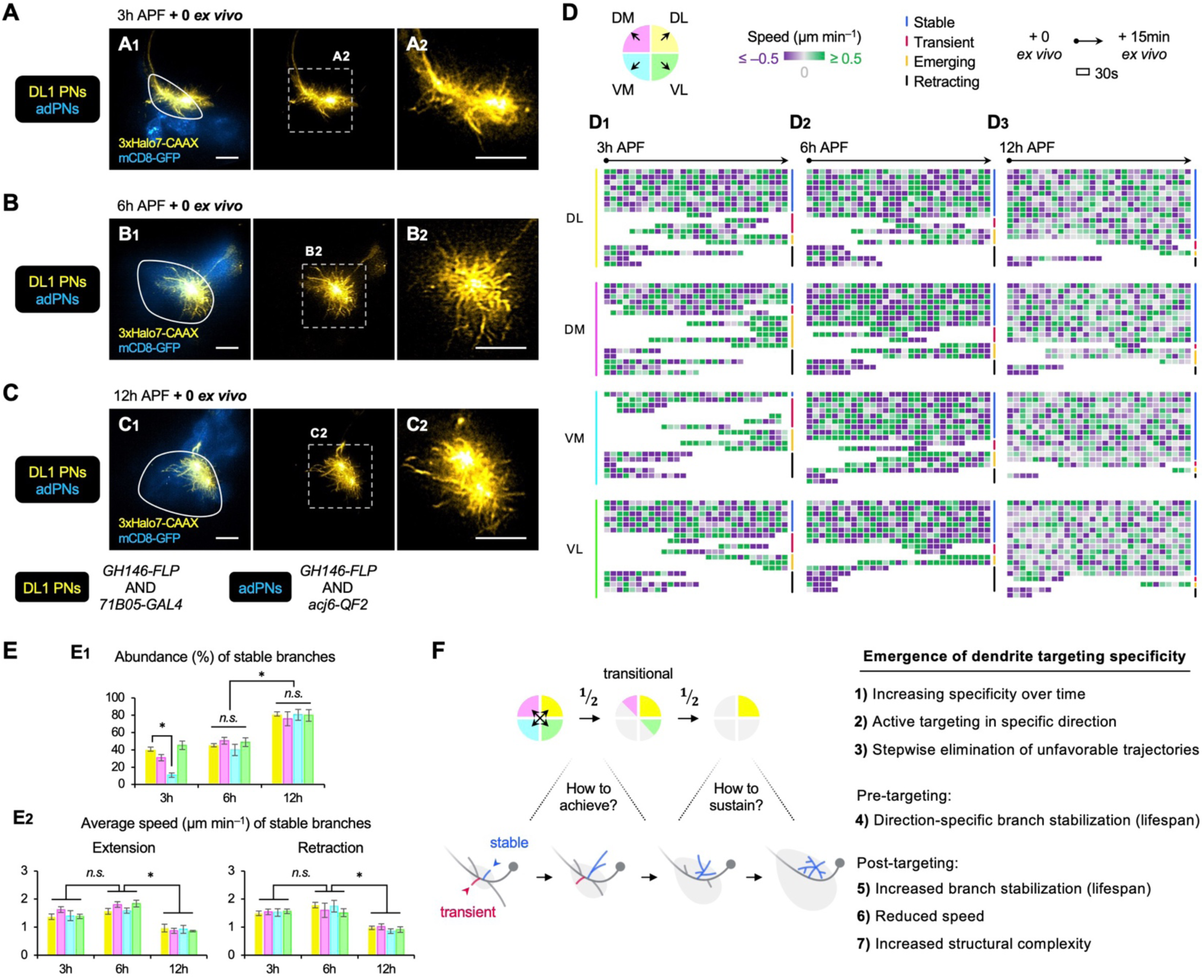
AO-LLSM time-lapse imaging reveals cellular mechanisms of dendrite targeting specificity. **(A–C)** AO-LLSM imaging of DL1 PNs (*71B05*+; labeled in yellow) and adPNs (*acj6*+; labeled in blue) in cultured brains dissected at 3h (**A**), 6h (**B**), and 12h (**C**) APF. Zoom-in, single z-section images of **A_1_**, **B_1_**, and **C_1_** (outlined in dashed boxes) are shown in **A_2_**, **B_2_**, and **C_2_**, respectively. **(D)** Single dendritic branch dynamics of DL1 PN at 3h (**D_1_**), 6h (**D_2_**), and 12h (**D_3_**) shown in Figure 7A**–C**. Terminal branches are analyzed and categorized based on the directions in which they extend. Their speeds are color-coded using dual purple-grey-green gradients (negative speeds, retraction; positive speeds, extension). Individual branches are also assigned into four categories: stable, transient, emerging, and retracting (color coded on the right; see Figure 7 **– figure supplement 1A**). Each block represents a 30s-window. Each row represents individual branch dynamics across the 15-min imaging period. **(E)** Quantification of the abundance (in percentage) of DL1 PN stable branches in indicated direction at 3h, 6h, and 12h (**E_1_**). Average speed of DL1 PN stable branches in indicated direction at 3h, 6h, and 12h (**E_2_**). DL1 PN sample size: 3h = 4; 6h = 3; 12h = 3. Error bars, SEM; *t* test; One-way ANOVA; *\**, *p* < 0.05; *n.s.*, *p* ≥ 0.05. SEM, standard error of the mean; *n.s.*, not significant. **(F)** Summary of mechanisms underlying the emergence of dendrite targeting specificity revealed by two-photon and AO-LLSM imaging of DL1 PN dendrites.

### AO-LLSM imaging suggests a cellular mechanism underlying dendrite targeting specificity

To capture fast dynamics of single dendritic branches, we performed dual-color adaptive optical lattice sheet microscopy (AO-LLSM) imaging (Chen et al., 2014; Wang et al., 2014; Liu et al., 2018) of PNs every 30 seconds for 15 minutes, following a protocol we recently established (Li et al., 2021; Li & Luo, 2021). We selected 3h, 6h, and 12h APF pupal brains double-labeled with DL1 PNs and bulk adPNs (**Figure 7A–C**; **Figure 7 – videos 1–3)**. The labeling of adPNs with GFP outlined PN cell bodies and the developing antennal lobe but not the degenerating one, presumably because the GFP in larval-specific dendrites was quickly quenched upon glial phagocytosis (Marin et al., 2005).

In the 15-min imaging window, we observed 4 types of terminal branches regardless of neuronal types or developmental stages: (1) stable branch that existed throughout the entire imaging time, (2) transient branch that was produced and eliminated within the imaging window, (3) emerging branch that was produced after imaging began, and (4) retracting branch that was eliminated within the imaging period (**Figure 7 – figure supplement 1A)**. To examine if terminal branch dynamics exhibit any directional preference, we assigned the branches according to their targeting directions (**Figure 7D)**. Extension and retraction events were defined when speed exceeded 0.5 μm/min. Terminal branches were selected for analyses as branches closer to the main process were too dense to resolve. **Figure 7D_1-3_** showed the dynamics of ∼15 randomly selected terminal branches in each direction from the representative 3h, 6h, and 12h APF DL1 PNs (**Figure 7A–C)**.

Quantitative analyses revealed that at 3h APF, DL1 PNs constantly produced, eliminated, extended, and retracted dendritic branches (**Figure 7A**, **Figure 7D_1_**, **Figure 7 – video 1)**. Even stable branches were not immobile. Rather, they spent comparable amounts of time extending and retracting at ∼1.5 μm/min (**Figure 7 – figure supplement 1A_1_, 1B)**. Transient, emerging, and retracting branches had similar, but more variable speed, ranging from 1 to 2.5 μm/min.

Although there was no correlation between targeting direction and frequency/speed of extension/retraction, the number of stable branches in the VM direction was significantly lower than in other directions across all 3h DL1 PN samples examined (**Figure 7E_1_)**. This suggests that even though dendritic branches were developed in every direction at the early stages, those branches in the VM direction were short-lived and might be eliminated by retraction. The direction-dependent stability/lifespan of dendritic branches on the timescale of seconds uncovered from AO-LLSM imaging explains why bulk dendrites in unfavorable trajectories failed to persist in long-term two-photon imaging.

From 6h to 12h APF, DL1 PNs no longer manifested direction-specific branch de/stabilization (**Figure 7B–C**, **Figure 7D_2–3_**, **Figure 7 – videos 2–3**). At the same developmental stage, stable branches in one direction appeared indistinguishable from those in other directions in terms of abundance, frequency, and speed (**Figure 7D_2–3_, Figure 7 – figure supplement 1C–D**). This suggests that the entire dendritic mass tends to stay in equilibrium upon arrival at target regions. At 12h APF, the abundance of stable branches of DL1 PNs was the highest (**Figure 7D–E_1_**). Also, the stable branches of 12h APF DL1 PNs moved at a significantly lower speed (∼1 μm/min) (**Figures 7E_2_**) and spent more time being stationary than those at 3h and 6h (**Figure 7 – figure supplement 1B–D**). The reduced branch dynamics at 12h APF is consistent with observations from two-photon imaging showing fewer bulk extension/retraction events in the final phase of targeting (**Figure 6E–F**). Despite the slowdown, dendritic arborization was evident in terminal branches of 12h APF DL1 PNs (**Figure 7 – figure supplement 1E**), indicating that PN dendrites are transitioning from simple to complex branch architectures. Although it remains unclear if there is a causal relationship between reduced branch dynamics and increased structural complexity, we propose that both contribute to the sustentation of dendrite targeting specificity.

In summary, AO-LLSM imaging reveals that PNs selectively stabilize branches in the direction towards the target and destabilize those in the opposite direction, providing a cellular basis of dendrite targeting specificity. Upon arrival at the target, the specificity is sustained through branch stabilization in a direction-independent manner (summarized in **Figure 7F_4–7_**).

### Embryonic-born PNs timely integrate into adult olfactory circuit by simultaneous dendritic pruning and re-extension

In earlier sections, we uncovered wiring logic of larval-born PN dendritic patterning and cellular mechanisms of dendrite targeting specificity used to initiate olfactory map formation (**Figures 3– 7)**. In this final section, we focused on embryonic-born PNs, which participate in both larval and adult olfactory circuits by reorganizing their processes (Marin et al., 2005). Our previous study demonstrates that embryonic-born PNs prune their larval-specific dendrites during early metamorphosis (Marin et al., 2005) (**Figure 1D_1–3_**). Here, we examined when and how embryonic-born PNs re-extend dendrites used in the adult olfactory circuit.

It is known that γ neurons of *Drosophila* mushroom body (γ Kenyon cells) and sensory Class IV dendritic arborization (C4da) neurons prune their processes between 4h and 18h APF and show no signs of re-extension at 18h APF (Lee et al., 2000; Watts et al., 2003; Lee et al., 2009). Do embryonic-born adPNs follow a similar timeframe? We first examined developing brains double-labeled for embryonic-born DA4l/VA6/VA2 adPNs (collectively referred to as ‘*lov+* PNs’) and early larval-born DC2 adPNs (**Figure 8A; Figure 8 – figure supplement 1**). We found that, by 12h APF, *lov+* PNs already sent adult-specific dendrites to a region ventromedial to DC2 PN dendrites (green arrowhead in **Figure 8A_3_**). This implies that *lov+* PNs have already caught up with DC2 PNs on dendrite development at this stage, and re-extension of *lov+* PN dendrites must have happened even earlier. Indeed, we observed *lov+* PN dendrites innervated the developing antennal lobe extensively at 6h APF (**Figure 8A_2_**). Such innervation was not observed at 0h APF (**Figure 8A_1_**). After 12h APF, the time course of *lov+* PN dendrite development was comparable to that of DC2 PNs (**Figure 8A_4–6_**).

**Figure 8.**
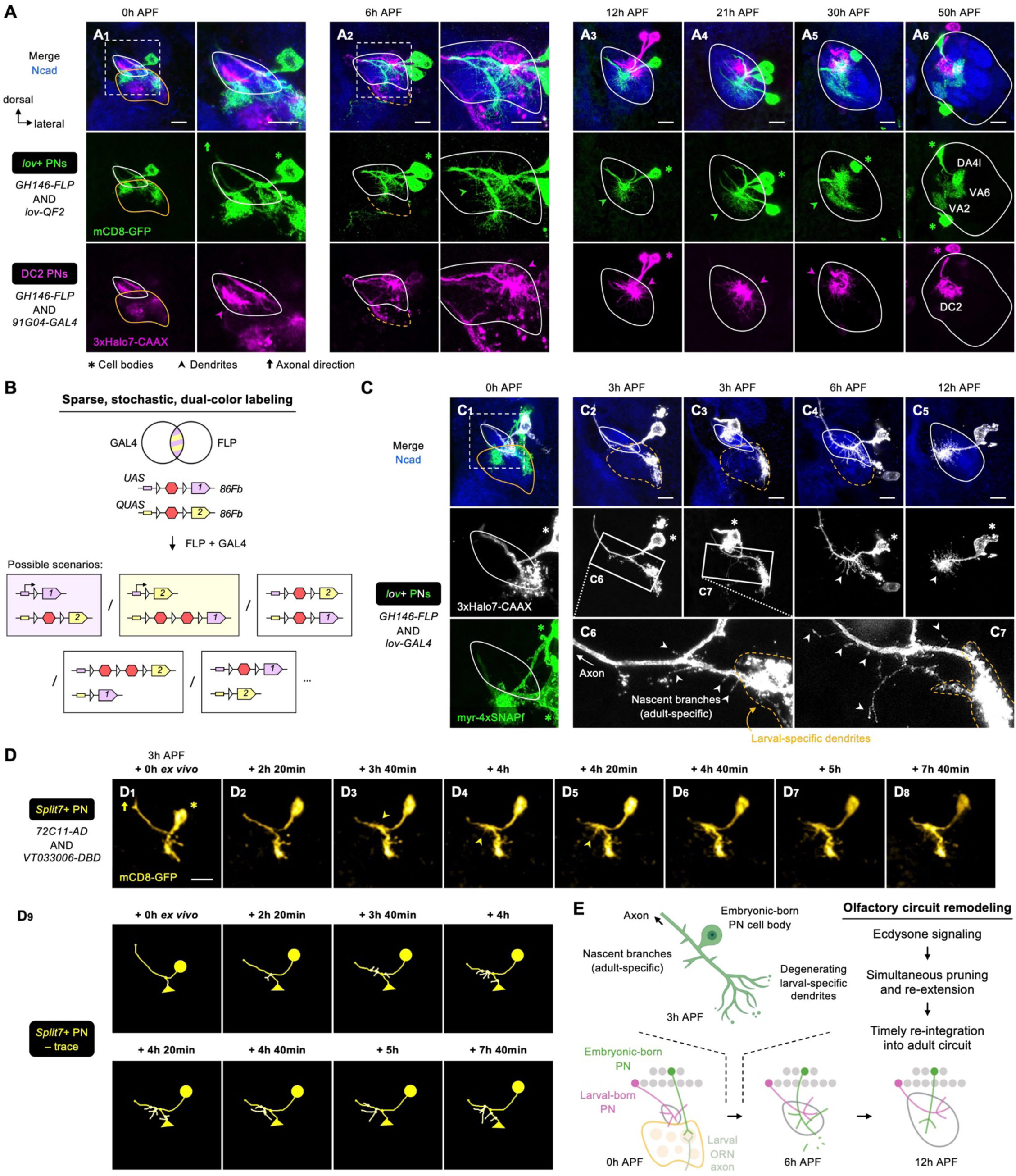
Embryonic-born PNs timely participate in olfactory map formation via simultaneous pruning and re-extension. **(A)** Confocal images of fixed brains at indicated stages showing dendrite development of *lov+* PNs (embryonic- born; labeled in green) and *91G04+* DC2 PNs (larval-born; labeled in magenta). As *91G04-GAL4* also labels some embryonic-born PNs from 0–6h APF, their processes are found in the larval-specific antennal lobe (**A_1, 2_**). Right columns of **A_1, 2_** show zoom-in of the dashed boxes. Green arrowhead in **A_2_** indicates robust dendrite re- extension of embryonic-born PNs across the developing antennal lobe at 6h APF. **(B)** Schematic of the sparse, stochastic, and dual-color labeling strategy. Following *FLP* and *GAL4* co- expression, random recombination of *FRT* sites yields expression of either reporter 1 or 2, or no labeling at all. **(C)** Sparse labeling of *lov*+ PNs (labeled in green; single-cell *lov+* PNs in grey) at indicated developmental stages. **C_6_** and **C_7_** are zoom-in images of the rectangular boxes in **C_2_** and **C_3_**, respectively. Arrowheads indicate nascent, adult-specific dendrites. Larval-specific dendrites are outlined by dashed orange lines. Arrows indicate axons projecting towards high brain centers. **(D)** Two-photon time-lapse imaging of a single embryonic-born PN (*Split7+*; pseudo-colored in yellow) in a brain dissected at 3h APF and cultured for 23h *ex vivo*. Arrowhead in **D_3_** denote the thickening of the main process. Arrowheads in **D_4, 5_** denote dendritic protrusions dorsal to larval-specific dendrites. **D_9_** show neurite tracing of the embryonic-born PN. Triangles in **D_9_** indicate the degenerating larval-specific dendrites. **(E)** Schematic summary of remodeling of embryonic-born PN dendrites. Following simultaneous pruning and re-extension, embryonic-born PNs timely integrates into adult olfactory circuit and, together with larval-born PNs, participate in the prototypic map formation.

To characterize dendritic re-extension at single-cell resolution, we developed a sparse, stochastic labeling strategy to label single *lov+* PNs (**Figure 8B**). We found that *lov+* PNs produced nascent branches from the main process dorsal to larval-specific dendrites as early as 3h APF (**Figure 8C_2–3_;** arrowheads in **Figure 8C_6–7_**). At 6h APF, when larval-specific dendrites were completely segregated from *lov+* PNs, robust extension of adult-specific dendrites was seen across the developing antennal lobe (**Figure 8C_4_**). These data indicate that *lov+* PNs re-extend their adult-specific dendrites at a more dorsal location before the larval-specific dendrites are completely pruned.

Do other embryonic-born PNs prune and re-extend their dendrites simultaneously? Like *lov* drivers, *Mz612-GAL4* labels embryonic-born PNs, one of which is VA6 PN (Marin et al., 2005). In 3h APF brains co-labeled for *Mz612+* and *lov+* PNs, we could unambiguously access 3 single embryonic-born PN types: (1) *lov+ Mz612–* PN, (2) *lov– Mz612*+ PN, and (3) *lov+ Mz612*+ PN (**Figure 8 – figure supplement 2A–B**). Tracing of individual dendritic branches showed that all these PNs already re-extended dendrites to varying extents prior to separation of larval-specific dendrites from the rest of the processes (**Figure 8 – figure supplement 2C**). Thus, concurrent pruning and re-extension apply to multiple embryonic-born PN types.

To capture the remodeling at higher temporal resolution, we performed two-photon time-lapse imaging of single embryonic-born PNs labeled by *Split7-GAL4* (**Figure 8D**, **Figure 8 – video 1, Figure 8 – figure supplement 3**). This *GAL4* labels one embryonic-born PN (either VA6 or VA2 PN) at early pupal stages but eight PN types at 24h APF (Xie et al., 2021). Initially (3h APF + 0h *ex vivo*), no adult-specific dendrites were detected in live *Split7+* PNs (**Figure 8D_1_**). The following ∼3h *ex vivo* saw thickening of the main process (arrowhead in **Figure 8D_3_**). From 4h *ex vivo* onwards, re-extension occurred in the presumed developing antennal lobe located dorsal to larval-specific dendrites (arrowheads in **Figure 8D_4–8_**; see traces in **Figure 8D_9_**). Live imaging of *Split7+* PNs also revealed that fragmentation of larval-specific dendrites occurred at the distal ends (**Figure 8 – figure supplement 3B_1–5_**), and the process leading to larval-specific dendrites gradually disappeared as pruning approached completion (**Figure 8 – figure supplement 3B_6–10_**). These observations suggest that pruning of embryonic-born PN dendrites is not initiated by severing at the proximal end. Distal-to-proximal pruning, rather than in the reversed direction, further supports concurrent but spatially segregated pruning and re-extension processes.

It has been shown that dendritic pruning of embryonic-born PNs requires ecdysone signaling in a cell-autonomous manner (Marin et al., 2005). We asked if the re-extension process also depends on ecdysone signaling. We expressed a dominant negative form of ecdysone receptor (EcR-DN) in most PNs (including *lov+* PNs) and monitored the development of *lov+* PN dendrites (**Figure 8 – figure supplement 4**). We found that inhibition of ecdysone signaling by *EcR-DN* expression not only suppressed pruning, but also blocked re-extension. This is consistent with a previous study reporting the dual requirement of ecdysone signaling in the pruning and re-extension of *Drosophila* anterior paired lateral (APL) neurons, although, unlike embryonic-born PNs, APL neurons prune and re-extend processes sequentially (at 6h and 18h APF, respectively) (Mayseless et al., 2018). We currently could not distinguish if the lack of re-extension is due to defective pruning, or if ecdysone signaling controls pruning and re-extension independently.

Taken together, our data demonstrate that embryonic-born PNs prune and re-extend dendrites simultaneously at spatially distinct regions, and that both processes require ecdysone signaling (**Figure 8E**). Such a ‘multi-tasking’ ability explains how embryonic-born PNs can re-integrate into the adult olfactory circuit and engage in its prototypic map formation in a timely manner.

## DISCUSSION

### Wiring logic for the prototypic olfactory map

Prior to this study, no apparent logic linking PN lineage, birth order, and adult glomerular position has been found. Our systematic analyses of dendritic patterning at the resolution of specific PN types across development identified wiring logic underlying the spatial organization of the prototypic olfactory map (**Figures 3 and 4**).

We found that PNs of a given lineage are grouped by birth order, and PNs of the same group share similar dendrite targeting specificity and timing. The strong correlation found between birth order and transcriptomic similarity among 0h APF adPNs (Xie et al., 2021) provides a molecular basis for executing this logic. In addition, we found that dendrites of adPNs and lPNs respectively pattern the antennal lobe in rotating and binary manners following birth order. Certain lineage-specific transcription factors are known to instruct dendrite targeting (Komiyama et al., 2003; Komiyama & Luo, 2007; Li et al., 2017; Xie et al., 2022), which may account for why the adPN and lPN dendritic maps differ. Cellular interactions among PN dendrites likely contribute further to the resulting olfactory map, given the robust dendritic dynamics seen during circuit formation (**Figures 5–8**). Our new tools for labeling and genetic manipulation of distinct PN types (**Figure 2**) will now enable in-depth investigations into the potential PN dendrite-dendrite interactions and molecular mechanisms leading to the initial map organization.

### Wiring logic evolves as development proceeds

After the initial map formation at 12h APF, dendrite positions in the antennal lobe could change substantially in the next 36 hours (for example, see DC2 PNs in **Figure 3B_4–6_** and DA1 and VA1d/DC3 PNs in **Figure 3C_4–7_**). These changes occur when dendrites of PNs with neighboring birth order begin to segregate and when ORN axons begin to invade the antennal lobe.

Accordingly, the oval-shaped antennal lobe turns into a globular shape (30–50h APF; **Figure 3C_6–7_**). These PN-autonomous and non-autonomous changes likely mask the initial wiring logic, explaining why previous studies, which mostly focused on examining the final glomerular targets in adults (Jefferis et al., 2001), have missed the earlier organization. Interestingly, the process of PN dendritic segregation coincides with the peak of PN transcriptomic diversity at 24h APF (Li et al., 2017; Xie et al., 2021).

Recent proteomics and genetic analyses have indicated that PN dendrite targeting is mediated by cell-surface proteins cooperating as a combinatorial code (Xie et al., 2022). The evolving wiring logic, which is consistent with the stepwise assembly of olfactory circuit (Hong & Luo, 2014), suggests the combinatorial codes are not static. We propose that PNs use a numerically simpler code for initial dendrite targeting (e.g., 18 adPN types being sorted into four groups with distinct targeting specificity; **Figure 3E**). Following the expansion of transcriptomic diversity, PNs acquire a more complex code mediating dendritic segregation of neighboring PNs and matching of PN dendrites and ORN axons. Functional characterization of differentially expressed genes between 12h and 24h APF PNs may provide molecular insights into how the degree of discreteness in the olfactory map arises.

### Selective branch stabilization as a cellular mechanism for dendrite targeting

Utilizing an early pupal brain explant culture system coupled with two-photon and AO-LLSM imaging (**Figure 5**), we presented the first time-lapse videos following dendrite development of a specific PN type – DL1 PNs (**Figures 6 and 7**). We found that DL1 PN dendrites initiate active targeting towards their dorsolateral target with direction-dependent branch stabilization. This directional selectivity provides a cellular basis of the emerging targeting specificity of PN dendrites at the beginning of olfactory map formation.

Although selective branch stabilization as a mechanism to achieve axon targeting specificity has been described in neurons in the vertebrate and invertebrate systems (e.g., Yates et al., 2001; Li et al., 2021), our time-lapse imaging showed, for the first time to our knowledge, that selective branch stabilization is also used to achieve dendrite targeting specificity.

Furthermore, AO-LLSM imaging revealed that selective stabilization and destabilization of dendritic branches occur on the timescale of seconds. As the rate of olfactory circuit development in the brain explants was slower than normal development (**Figure 5F**), we might have captured PN dendritic dynamics in slow motion. Using AO-LLSM for high spatiotemporal resolution imaging, we just begin to appreciate how fast PN dendrites are coordinating trajectory choices with branch stabilization to make the appropriate decision. Having characterized the dendritic branch dynamics of the wild-type DL1 PNs, we have set the stage for future studies addressing how positional cues and the downstream signaling instruct wiring, and whether other PN types follow similar rules as DL1 PNs.

### Simultaneous pruning and re-extension as novel remodeling mechanism for neuronal remodeling

Our data on embryonic-born adPN dendrite development reveals a novel mode of neuronal remodeling during metamorphosis (**Figure 8**). In mushroom body γ neurons and body wall somatosensory neurons, two well-characterized systems, larval-specific neurites are first pruned, followed by re-extension of adult-specific processes (Watts et al., 2003; Williams and Truman, 2005; Yaniv and Schuldiner, 2016). However, embryonic-born adPNs prune larval-specific dendrites and re-extend adult-specific dendrites simultaneously but at spatially separated regions. Such spatial segregation suggests that regional external cues could elicit compartmentalized downstream signals leading to opposite effects on the dendrites. Subcellular compartmentalization of signaling and cytoskeletal organization has been observed in diverse neuron types across species (Rolls et al., 2007; Kanamori et al., 2013; O’Hare et al., 2022).

Why do embryonic-born adPNs ‘rush’ to re-extend dendrites? During normal development, it takes at least 18 hours for embryonic-born adPNs to produce and properly target dendrites (growth at 3–6h APF, initial targeting at 6–12h APF, and segregation at 21–30h APF). Given that the dendritic re-extension of embryonic-born PNs is ecdysone dependent (**Figure 8 – figure supplement 4**), if the PNs did not re-extend dendrites at 3h APF, they would have to wait for the next ecdysone surge at ∼20h APF (Thummel, 2001), which might be too late for their dendrites to engage in the prototypic map formation. Thus, embryonic-born PNs develop a remodeling strategy that coordinates with the timing of systemic ecdysone release. By simultaneous pruning and re-extension, embryonic-born adPNs timely re-integrate into the adult prototypic map that readily serves as target for subsequent ORN axon innervation.

In conclusion, our study highlights the power and necessity of type-specific neuronal access and time-lapse imaging to identify wiring logic and mechanisms underlying the origin of an olfactory map. Applying similar approaches to other developing neural maps across species should broaden our understanding of the generic and specialized designs that give rise to functional maps with diverse architectures.

## MATERIALS AND METHODS

### *Drosophila* stocks and husbandry

Flies were maintained on standard cornmeal medium at 25°C. Fly lines used in this study included *GH146-FLP* (Hong et al., 2009), *QUAS-FRT-stop-FRT-mCD8-GFP* (Potter et al., 2010), *UAS-mCD8-GFP* (Lee & Luo, 1999), *UAS-mCD8-FRT-GFP-FRT-RFP* (Stork et al., 2014), *VT033006-GAL4* (Tirian & Dickson, 2017), *Mz19-GAL4* (Jefferis et al., 2004), *91G04- GAL4* (Jenett et al., 2012), *Mz612-GAL4* (Marin et al., 2005), *71B05-GAL4* (Jenett et al., 2012), *Split7-GAL4* (Xie et al., 2021), *QUAS-FLP* (Potter et al., 2010) and *UAS-EcR.B1-ΔC655.F645A* (Cherbas et al., 2003). The following GAL4 lines were obtained from Bloomington *Drosophila* Stock Center (BDSC): *tsh-GAL4* (BDSC #3040) and *lov-GAL4* (BDSC #3737).

The following two stocks were used for MARCM analyses: (1) *UAS-mCD8-GFP, hs- FLP; FRT^G13^, tub-GAL80;; GH146-GAL4*, and (2) *FRT^G13^, UAS-mCD8-GFP* (Lee & Luo, 1999).

The following lines were generated in this study: *UAS-FRT^10^-stop-FRT^10^-3xHalo7-CAAX* (on either II or III chromosome), *UAS-FRT-myr-4xSNAPf-FRT-3xHalo7-CAAX* (III), *UAS-FRT-myr-mGreenLantern-FRT-3xHalo7-CAAX* (II), *QUAS-FRT-stop-FRT-myr-4xSNAPf* (III), *run- T2A-FLP* (X), *acj6-T2A-FLP* (X), *acj6-T2A-QF2* (X), *CG14322-T2A-QF2* (III) and *lov-T2A- QF2* (II).

### Drosophila genotypes

**Figure 1D**, **Figure 1 – figure supplement 1, Figure 1 – figure supplement 2**: *run-T2A-FLP/+; UAS-mCD8-FRT-GFP-FRT-RFP/+; VT033006-GAL4/+*

**Figure 3A**: *acj6-T2A-QF2/+; GH146-FLP, QUAS-FRT-stop-FRT-mCD8-GFP/UAS-FRT^10^- stop-FRT^10^-3xHalo7-CAAX; 71B05-GAL4/+*

**Figure 3B**, **Figure 3 – figure supplement 1C**: *GH146-FLP/UAS-FRT^10^-stop-FRT^10^-3xHalo7- CAAX; 91G04-GAL4/CG14322-T2A-QF2, QUAS-FRT-stop-FRT-myr-4xSNAPf*

**Figure 3C**: *acj6-T2A-FLP/+; Mz19-GAL4; UAS-FRT-myr-4xSNAPf-FRT-3xHalo7-CAAX/+*

**Figure 3D**, **Figure 3 – figure supplement 2, Figure 3 – figure supplement 3**: *UAS-mCD8- GFP, hs-FLP/+; FRT^G13^, tub-GAL80/FRT^G13^, UAS-mCD8-GFP;; GH146-GAL4 (IV)/+*

**Figure 3 – figure supplement 1A**: *GH146-FLP/UAS-FRT^10^-stop-FRT^10^-3xHalo7-CAAX; 71B05-GAL4/+*

**Figure 3 – figure supplement 1B**: *GH146-FLP/UAS-FRT^10^-stop-FRT^10^-3xHalo7-CAAX; 91G04-GAL4/+*

**Figure 4A**, **Figure 4 – figure supplement 1**: *GH146-FLP, UAS-FRT^10^-stop-FRT^10^-3xHalo7- CAAX/tsh-GAL4; CG14322-T2A-QF2, QUAS-FRT-stop-FRT-myr-4xSNAPf/+*

**Figure 4B**: *UAS-mCD8-GFP, hs-FLP/+; FRT^G13^, tub-GAL80/FRT^G13^, UAS-mCD8-GFP;; GH146-GAL4 (IV)/+*

**Figure 4 – figure supplement 2**: *acj6-T2A-FLP/+; tsh-GAL4, UAS-mCD8-FRT-GFP-FRT-RFP* **Figure 5E**, **Figure 5 – video 1**: *run-T2A-FLP/+; UAS-FRT-myr-mGreenLantern-FRT-3xHalo7- CAAX/+; VT033006-GAL4/+*

**Figure 5F**: *UAS-mCD8-GFP/+; VT033006-GAL4/+*

**Figure 5G1**: *GH146-FLP/UAS-FRT^10^-stop-FRT^10^-3xHalo7-CAAX; 71B05-GAL4/+*

**Figure 5G2**: *GH146-FLP/tsh-GAL4; UAS-FRT^10^-stop-FRT^10^-3xHalo7-CAAX/+*

**Figure 5 – figure supplement 1, Figure 5 – video 2**: *acj6-T2A-FLP/+; Mz19-GAL4/UAS-FRT- myr-mGreenLantern-FRT-3xHalo7-CAAX*

**Figure 6A**, **Figure 6 – figure supplement 1, Figure 6 – video 1**: *UAS-mCD8-GFP, hs-FLP/+; FRT^G13^, tub-GAL80/FRT^G13^, UAS-mCD8-GFP;; GH146-GAL4 (IV)/+*

**Figure 7A–C**, **Figure 7 – figure supplement 1, Figure 7 – videos 1–3**: *acj6-T2A-QF2/+; GH146-FLP, QUAS-FRT-stop-FRT-mCD8-GFP/UAS-FRT^10^-stop-FRT^10^-3xHalo7-CAAX; 71B05-GAL4/+*

**Figure 8A**, **Figure 8 – figure supplement 1**: *GH146-FLP, QUAS-FRT-stop-FRT-mCD8- GFP/lov-T2A-QF2; UAS-FRT^10^-stop-FRT^10^-3xHalo7-CAAX/91G04-GAL4*

**Figure 8C**: *GH146-FLP/lov-GAL4; UAS-FRT^10^-stop-FRT^10^-3xHalo7-CAAX/QUAS-FRT-stop-FRT-myr-4xSNAPf*

**Figure 8D**, **Figure 8 – figure supplement 3, Figure 8 – video 1**: UAS-mCD8-GFP/+; Split7- GAL4 (i.e. FlyLight SS01867: 72C11-p65ADZp; VT033006-ZpGDBD)/+

**Figure 8 – figure supplement 2**: *GH146-FLP, QUAS-FRT-stop-FRT-mCD8-GFP/lov-T2A- QF2, Mz612-GAL4; UAS-FRT^10^-stop-FRT^10^-3xHalo7-CAAX/+*

**Figure 8 – figure supplement 4A**: *lov-T2A-QF2, QUAS-FLP/+; VT033006-GAL4/UAS-mCD8- FRT-GFP-FRT-RFP*

**Figure 8 – figure supplement 4B**: *lov-T2A-QF2, QUAS-FLP/UAS-EcR-DN; VT033006- GAL4/UAS-mCD8-FRT-GFP-FRT-RFP*

### MARCM clonal analyses

MARCM clonal analyses have been previously described (Lee & Luo, 1999). Larvae of the genotype *UAS-mCD8-GFP, hs-FLP/+; FRT^G13^, tub-GAL80/FRT^G13^, UAS-mCD8-GFP;; GH146-GAL4/+* were heat shocked at 37°C for 1 hour. To label the first-born DL1 PNs, heat shock was applied at <24h after larval hatching (ALH). MARCM clones of early, middle and late larval- born PNs were generated by applying heat shocks at 48h, 72h and >96h ALH, respectively. As larvae developed at different rates (Tennessen & Thummel, 2011), we reasoned that even if we could collect 0h–2h ALH larvae, their development might have varied by the time of heat shock. To minimize the effects of unsynchronized development, we selected those heat-shocked larvae that were among the first to form puparia and collected these white pupae in a ∼3-hour window for the clonal analyses.

### Transcriptomic analyses

Transcriptomic analyses have been described previously (Xie et al., 2021). tSNE plots and dot plots were generated in Python using PN single-cell RNA sequencing data and code available at https://github.com/Qijing-Xie/FlyPN_development.

### Generation of *T2A-QF2/FLP* lines

To generate a *T2A-QF2/FLP* donor vector for *acj6* (we used the same strategy for *run, CG14322* and *lov*), a ∼2000-bp genomic sequence flanking the stop codon of *acj6* was PCR amplified and introduced into *pCR-Blunt II-TOPO* (ThermoFisher Scientific #450245), forming *pTOPO-acj6*. To build *pTopo-acj6-T2A-QF2*, *T2A-QF2* including *loxP*-flanked *3xP3-RFP* was PCR amplified from *pBPGUw-HACK-QF2* (Addgene #80276), followed by insertion into *pTOPO-acj6* right before the stop codon of *acj6* by DNA assembly (New England BioLabs #E2621S). To generate *T2A-FLP*, we PCR amplified *FLP* from genomic DNA of *GH146-FLP* strain. *QF2* in *pTopo-acj6-T2A-QF2* was then replaced by *FLP* through DNA assembly. Using CRISPR Optimal Target Finder (Gratz et al., 2014), we selected a 20-bp gRNA target sequence that flanked the stop codon and cloned it into *pU6-BbsI-chiRNA* (Addgene #45946). If the gRNA sequence did not flank the stop codon, silent mutations were introduced at the PAM site of the donor vector by site-directed mutagenesis. Donor and gRNA vectors were co-injected into *Cas9* embryos in-house or through BestGene.

### Generation of FLP-out reporters

To generate *pUAS-FRT^10^-stop-FRT^10^-3xHalo7-CAAX*, *FRT^10^-stop-FRT^10^* was PCR amplified from *pUAS-FRT^10^-stop-FRT^10^-mCD8-GFP* (Li et al., 2021) and inserted into *pUAS-3xHalo7- CAAX* (Addgene #87646) through NotI and DNA assembly.

To generate pUAS-FRT-myr-4xSNAPf-FRT-3xHalo7-CAAX, we first PCR amplified myr- 4xSNAPf from pUAS-myr-4xSNAPf (Addgene #87637) using FRT-containing primers. FRT-myr- 4xSNAPf-FRT was then introduced into pCR-Blunt II-TOPO, forming pTOPO-FRT-myr- 4xSNAPf-FRT. Using NotI-containing primers, FRT-myr-4xSNAPf-FRT was PCR amplified and subcloned into pUAS-3xHalo7-CAAX through NotI.

To generate pUAS-FRT-myr-mGreenLantern-FRT-3xHalo7-CAAX, we first PCR amplified mGreenLantern from pcDNA3.1-mGreenLantern (Addgene #161912). Using MluI and XbaI, we replaced 4xSNAPf in pUAS-myr-4xSNAPf with mGreenLantern to build pUAS-myr-mGreenLantern. myr-mGreenLantern was PCR amplified with the introduction of FRT sequence, followed by insertion into pCR-Blunt II-TOPO. Using the NotI-containing primers, FRT-myr-mGreenLantern-FRT was PCR amplified and subcloned into pUAS-3xHalo7-CAAX through NotI.

To generate pQUAS-FRT-stop-FRT-myr-4xSNAPf, we first PCR amplified FRT-stop from pJFRC7-20XUAS-FRT-stop-FRT-mCD8-GFP (Li et al., 2021) and inserted it into pTOPO- FRT-myr-4xSNAPf-FRT through DNA assembly to form pTOPO-FRT-stop-FRT-myr-4xSNAPf- FRT. Using NotI-containing forward and KpnI-containing reverse primers, FRT-stop-FRT-myr- 4xSNAPf was PCR amplified and subcloned into p10XQUAST. p10XQUAST was generated using p5XQUAS (Addgene #24349) and p10xQUAS-CsChrimson (Addgene #163629).

*attP24* and *86Fb* landing sites were used for site-directed integration.

### Immunofluorescence staining and confocal imaging

Fly brain dissection for immunostaining and live imaging has been described (Wu & Luo, 2006). Briefly, brains were dissected in phosphate buffered saline (PBS) and fixed with 4% paraformaldehyde in PBS for 20 minutes on a nutator at room temperature. Fixed brains were washed with 0.1% Triton X-100 in PBS (PBST) for 10 min twice. After blocking with 5% normal donkey serum in PBST for 1 hour at room temperature, the brains were incubated with primary antibodies overnight at 4°C. After PBST wash, brains were incubated with secondary antibodies (1:1000; Jackson ImmunoResearch) in dark for 2 hours at room temperature. Washed and mounted brains were imaged with confocal laser scanning microscopy (ZEISS LSM 780; LSM 900 with Airyscan 2). Images were processed with ImageJ. Neurite tracing images were generated using Simple Neurite Tracer (SNT) (Arshadi et al., 2021). Primary antibodies used included chicken anti-GFP (1:1000; Aves Lab #GFP-1020), rabbit anti-DsRed (1:500; TaKaRa #632496), rat anti-Cadherin DN (1:30; Developmental Studies Hybridoma Bank DSHB DN-Ex#8 supernatant), and mouse anti-Bruchpilot (1:30; DSHB nc82 supernatant).

### Chemical labeling

Chemical labeling of *Drosophila* brains has been described (Kohl et al., 2014). Janelia Fluor (JF) Halo and SNAP ligands (stocks at 1 mM) were gifts from Dr. Luke Lavis (Grimm et al., 2017, 2021).

Fixed brains were washed with PBST for 5 min, followed by incubation with Halo and/or SNAP ligands (diluted in PBS) for 45 min at room temperature. Brains were then washed with PBST for 5 min, followed by blocking and immunostaining if necessary. For co-incubation of Halo and SNAP ligands, JF503-cpSNAP (1:1000) and JF646-Halo (1:1000) were used.

Alternatively, JFX650-SNAP (1:1000) and JFX554-Halo (1:10000) were used. When only Halo ligands were needed, either JF646-Halo or JF635-Halo (1:1000) was used.

For live brain imaging, dissected brains were incubated with Halo ligands diluted in culture media (described below) for 30 min at room temperature. For two-photon imaging, JF570-Halo was used at 1:5000. For AO-LLSM imaging, following JF646-Halo incubation at 1:1000, the brains were incubated with 1 µM Sulforhodamine 101 (Sigma) for 5 min at room temperature. The brains were then briefly washed with culture media before imaging.

### Brain explant culture setup and medium preparation

Brain explant culture setup was modified based on Li et al., 2021; Li & Luo, 2021. A Sylgard plate with a thickness of ∼ 2 millimeters was prepared by mixing base and curing agent at 10:1 ratio (DOW SYLGARD^TM^ 184 Silicone Elastomer Kit). The mixture was poured into a 60 mm x 15 mm dish in which it was cured for two days at room temperature. Once cured, the plate was cut into small squares (∼15 mm x ∼15 mm). Indentations were created based on the size of an early pupal brain using a No.11 scalpel. Additional slits were made around the indentations for attaching imaginal discs which served as anchors to hold the brain position. A square Sylgard piece was then placed in a 60 mm x 15 mm dish or on a 25-mm round coverslip in preparation for two-photon/AO-LLSM imaging.

Culture medium was prepared based on published methods (Rabinovich et al., 2015; Li and Luo, 2021; Li et al., 2021). The medium contained Schneider’s *Drosophila* Medium (ThermoFisher Scientific #21720001), 10% heat-inactivated Fetal Bovine Serum (ThermoFisher Scientific #16140071), 10 µg/mL human recombinant insulin (ThermoFisher Scientific #12585014; stock = 4 mg/mL), 1:100 Penicillin-Streptomycin (ThermoFisher Scientific #15140122). For 0h–6h APF brain culture, 0.5 mM ascorbic acid (Sigma #A4544; stock concentration = 50 mg/mL in water) was included. 20-hydroxyecdysone (Sigma #H5142; stock concentration = 1 mg/mL in ethanol) was used for 0h–6h and 12h brain explants at 20 µM and 2 µM, respectively. Culture medium was oxygenated for 20 minutes before use.

### Single- and dual-color imaging with two-photon microscopy

Single- and dual-color imaging of PNs were performed at room temperature using a custom-built two-photon microscope (Prairie Technologies) with a Chameleon Ti:Sapphire laser (Coherent) and a 16X water-immersion objective (0.8 NA; Nikon). Excitation wavelength was set at 920 nm for GFP imaging, and at 935 nm for co-imaging of mGreenLantern and JF570-Halo. *z*-stacks were obtained at 4-µm increments (10-µm increments for **Figure 5 – video 1**). Images were acquired at a resolution 1024 x 1024 pixel^2^ (512 x 512 for **Figure 5 – video 1**), with a pixel dwell time of 6.8 µs and an optical zoom of 2.1, and at a frequency every 20 minutes for 8–23 hours.

### Dual-color imaging with AO-LLSM

For AO-LLSM based imaging, the excitation and detection objectives along with the 25-mm coverslip were immersed in ∼40 mL of culture medium at room temperature. Explant brains held on Sylgard plate were excited simultaneously using 488 nm (for GFP) and 642 nm (for JF-646) lasers operating with ∼2–10 mW input power to the microscope (corresponding to ∼10–50 µW at the back aperture of the excitation objective). An exposure time of 20–50 msec was used to balance imaging speed and signal-to-noise ratio (SNR). Dithered lattice light-sheet patterns with an inner/outer numerical aperture of 0.35/0.4 or 0.38/0.4 were used. The optical sections were collected by an axial step size of 250 nm in the detection objective coordinate, with a total of 81– 201 steps (corresponding to a total axial scan range of 20–50 µm). Emission light from GFP and JF-646 was separated by a dichromatic mirror (Di03-R561, Semrock, IDEX Health & Science, LLC, Rochester, NY) and captured by two Hamamatsu ORCA-Fusion sCMOS cameras simultaneously (Hamamatsu Photonics, Hamamatsu City, Japan). Prior to the acquisition of the time series data, the imaged volume was corrected for optical aberrations using two-photon guide star based adaptive optics method (Chen et al., 2014; Wang et al., 2014; Liu et al., 2018). Each imaged volume was deconvolved using Richardson-Lucy algorithm on HHMI Janelia Research Campus’ or Advanced Bioimaging Center’s computing cluster (https://github.com/scopetools/cudadecon, https://github.com/abcucberkeley/LLSM3DTools) with experimentally measured point spread functions obtained from 100 or 200 nm fluorescent beads (Invitrogen FluoSpheres^TM^ Carboxylate-Modified Microspheres, 505/515 nm, F8803, FF8811). The AO-LLSM was operated using a custom LabVIEW software (National Instruments, Woburn, MA).

### Statistics

For data analyses, t-test and one-way ANOVA were used to determine *p* values as indicated in the figure legend for each graph, and graphs were generated using Excel. Exact *p* values were provided in Source Data files.

### Material and data availability

All reagents generated in this study are available from the lead corresponding author without restriction. Figure 3 - Source Data 1, Figure 5 - Source Data 1, Figure 6 - Source Data 1, and Figure 7 - Source Data 1 contain the numerical and statistical data used to generate the figures.

## ACKNOWLEDGEMENTS

We thank the Luo lab members for constructive feedback on the manuscript; Tzumin Lee for sharing equipment at Janelia Research Campus; Luke Lavis for sharing JF dyes. E.B. and L.L. are HHMI investigators. This work was support by a grant from NIH (R01 DC005982 to L.L.). GL and S.U. are funded by Philomathia Foundation. S.U. is funded by Chan Zuckerberg Initiative Imaging Scientist program. S.U. is a Chan Zuckerberg Biohub Investigator. T.L. was supported by NIH 1K99DC01883001.

---

**Figure 1 – figure supplement 1.**
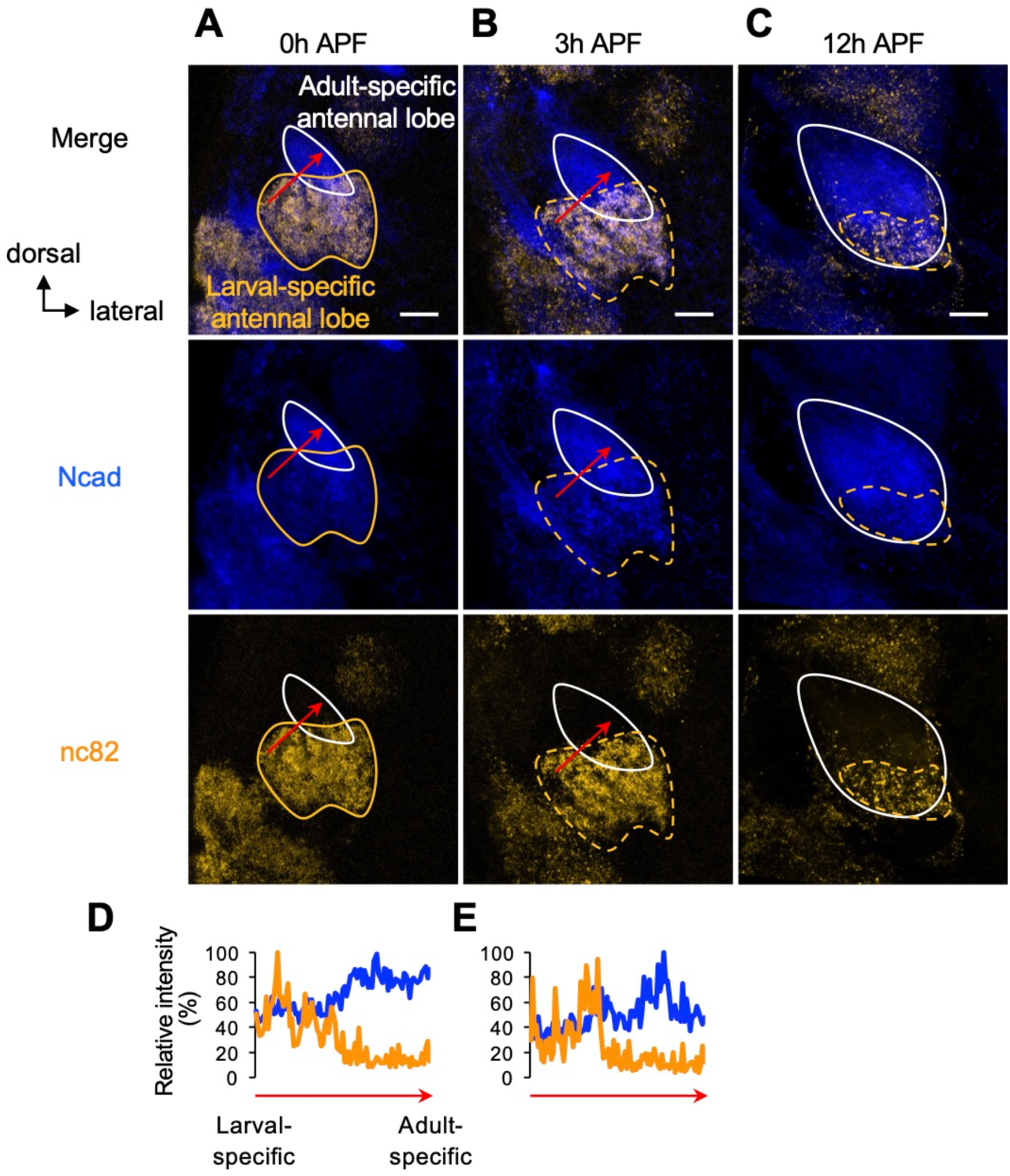
Visualization of larval- and adult-specific antennal lobes by co-staining of Ncad and nc82. **(A–C)** Ncad and nc82 staining of antennal lobes shown in **Figure 1D_1–3_**. Adult-specific antennal lobes characteristic of strong Ncad and weak nc82 staining are outlined by white solid line. Larval-specific antennal lobes characteristic of weak Ncad and strong nc82 are outlined by orange solid/dashed line. Note that larval-specific antennal lobes are more anterior to adult-specific antennal lobes, and thus appear overlapping in these z-projections. **(D, E)** Quantification of the relative intensity (%) of Ncad and nc82 staining from larval- to adult-specific antennal lobes at 0h APF **(D)** and 3h APF **(E)** (red arrows in **D** and **E** correspond to those in **A** and **B**, respectively). See Figure 1 legend for common notations.

**Figure 1 – figure supplement 2.**
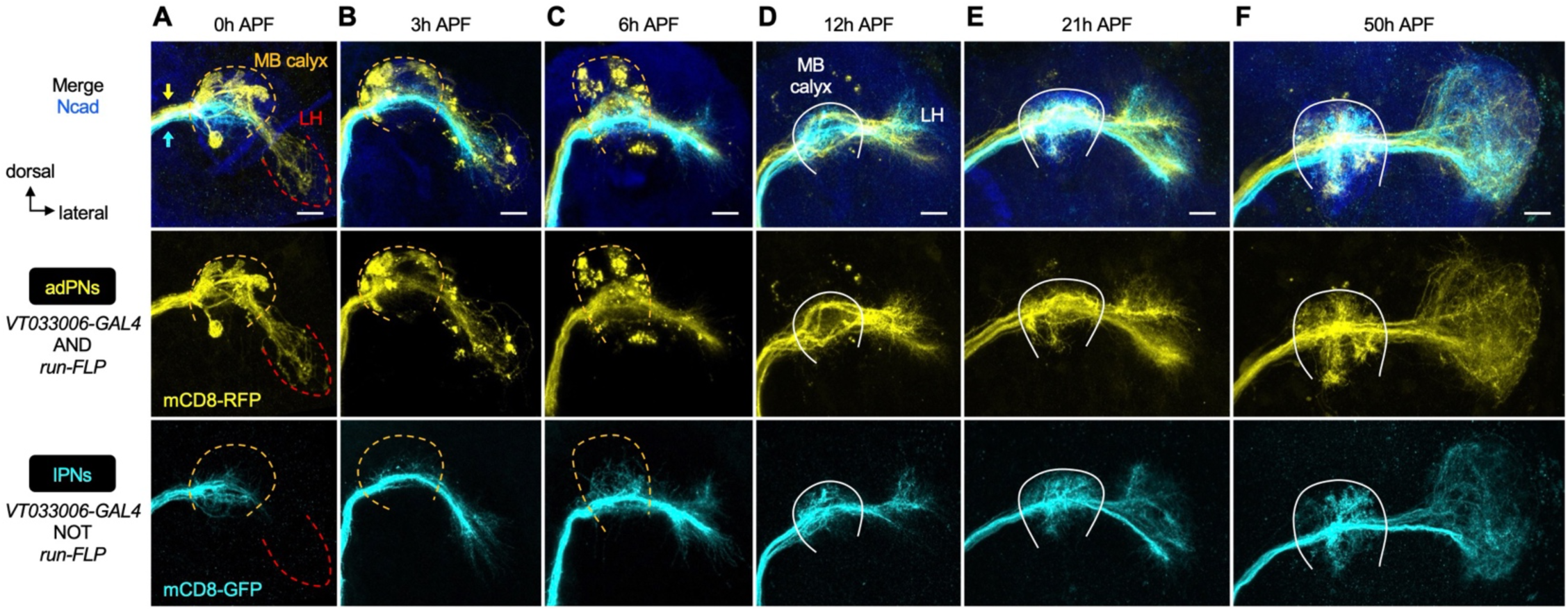
PN axon development across pupal stages. **(A–F)** Staining of fixed brains at indicated stages showing axon development of adPNs (*VT033006+ run+*; labeled in yellow) and lPNs (*VT033006+ run–*; labeled in cyan). Yellow and cyan arrows in **A** indicate the segregation of adPN and lPN axons along the inner antennocerebral tract. MB: mushroom body; LH: lateral horn. MB calyx (where PN axons and Kenyon cells of the mushroom body form synapses) and LH neuropils (where PN axons form synapses with their postsynaptic target neurons) are outlined as follows. In **A–C**, orange dashed line denotes the degeneration of larval-specific MB calyx. In **A**, larval-specific LH located more ventrally is outlined by red dashed line. In **D–F**, the developing adult-specific MB calyx is outlined by white solid line and adult-specific LH is to the right of the calyx. See Figure 1 legend for common notations.

**Figure 2 – figure supplement 1.**
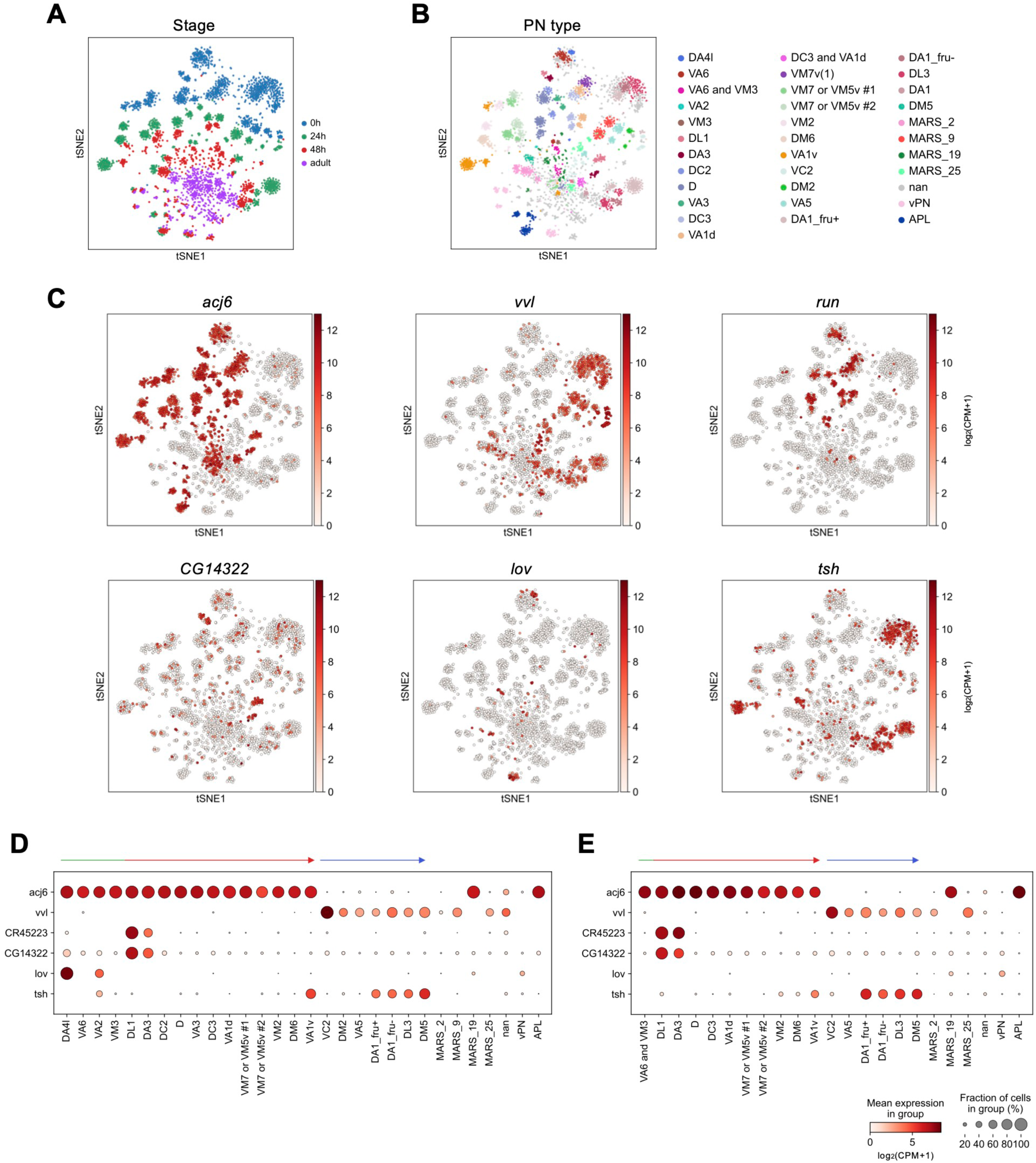
Expression of PN marker genes across development. **(A–C)** tSNE plots of PN transcriptomes, color-coded according to developmental stages (**A**), PN types (**B**), or the expression levels of indicated gene [log_2_(CPM+1)] (**C**) using scRNA-seq data from Xie et al., 2021. **(D, E)** Dot plot showing the expression levels of *acj6*, *vvl*, *CR45223*, *CG14322*, *lov*, and *tsh* in PNs [log_2_(CPM+1)] at 24h APF (**D**) and 48h APF (**E**).

**Figure 2 – figure supplement 2.**
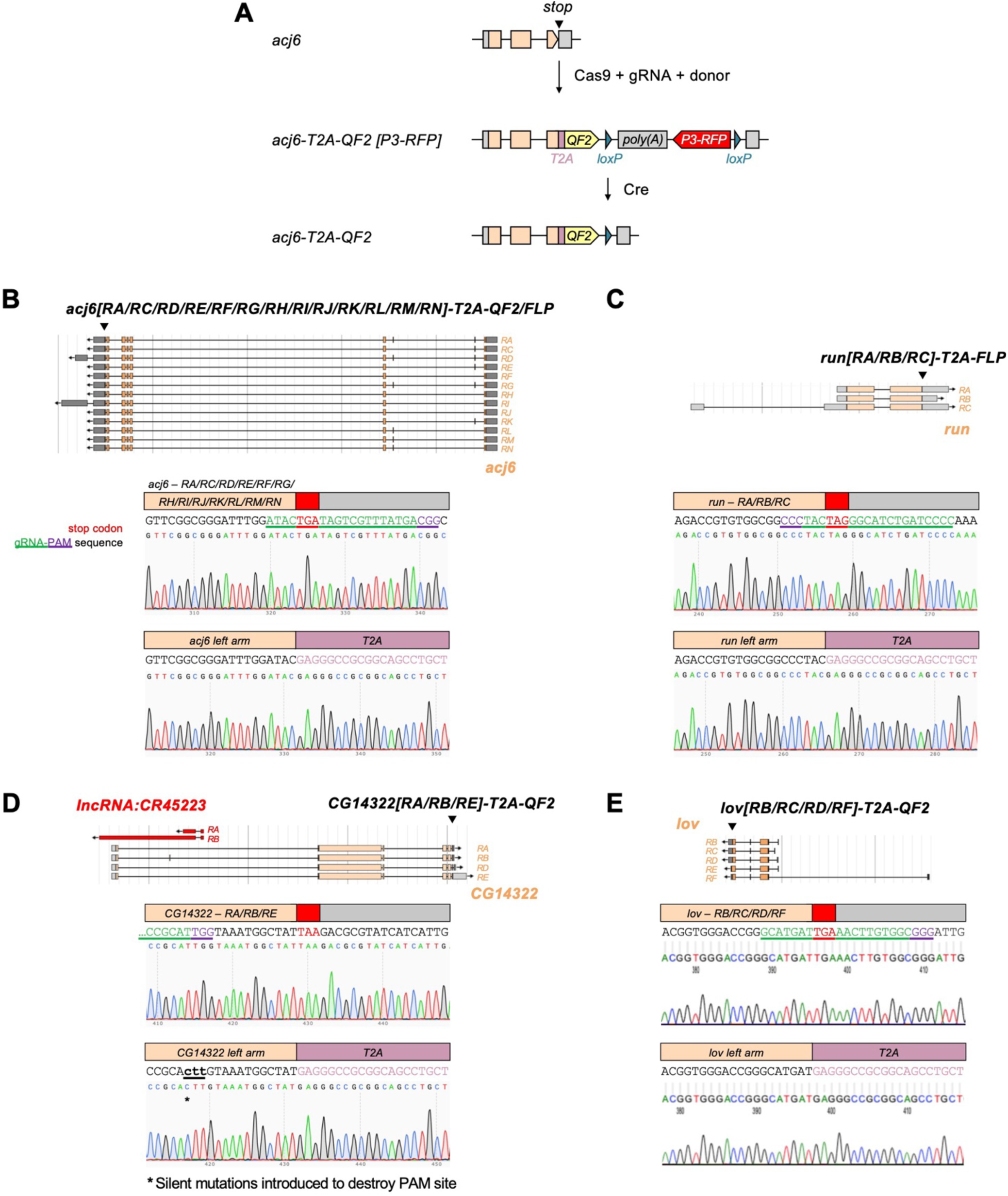
Generation of *T2A-QF2/FLP* transgenic flies by CRISPR/Cas9. **(A)** Schematic of generation of transgenic driver lines by CRISPR/Cas9. *acj6-T2A-QF2* is shown as an example. **(B–E) Top**: Transcripts of *acj6* (**B**), *run* (**C**), *CG14322* (**D**), and *lov* (**E**) visualized using FlyBase JBrowse. **Bottom**: Targeted insertion of *T2A-QF2/FLP* right before the stop codon of the endogenous gene. Stop codon and gRNA-PAM sequence are color-coded as indicated.

**Figure 2 – figure supplement 3.**
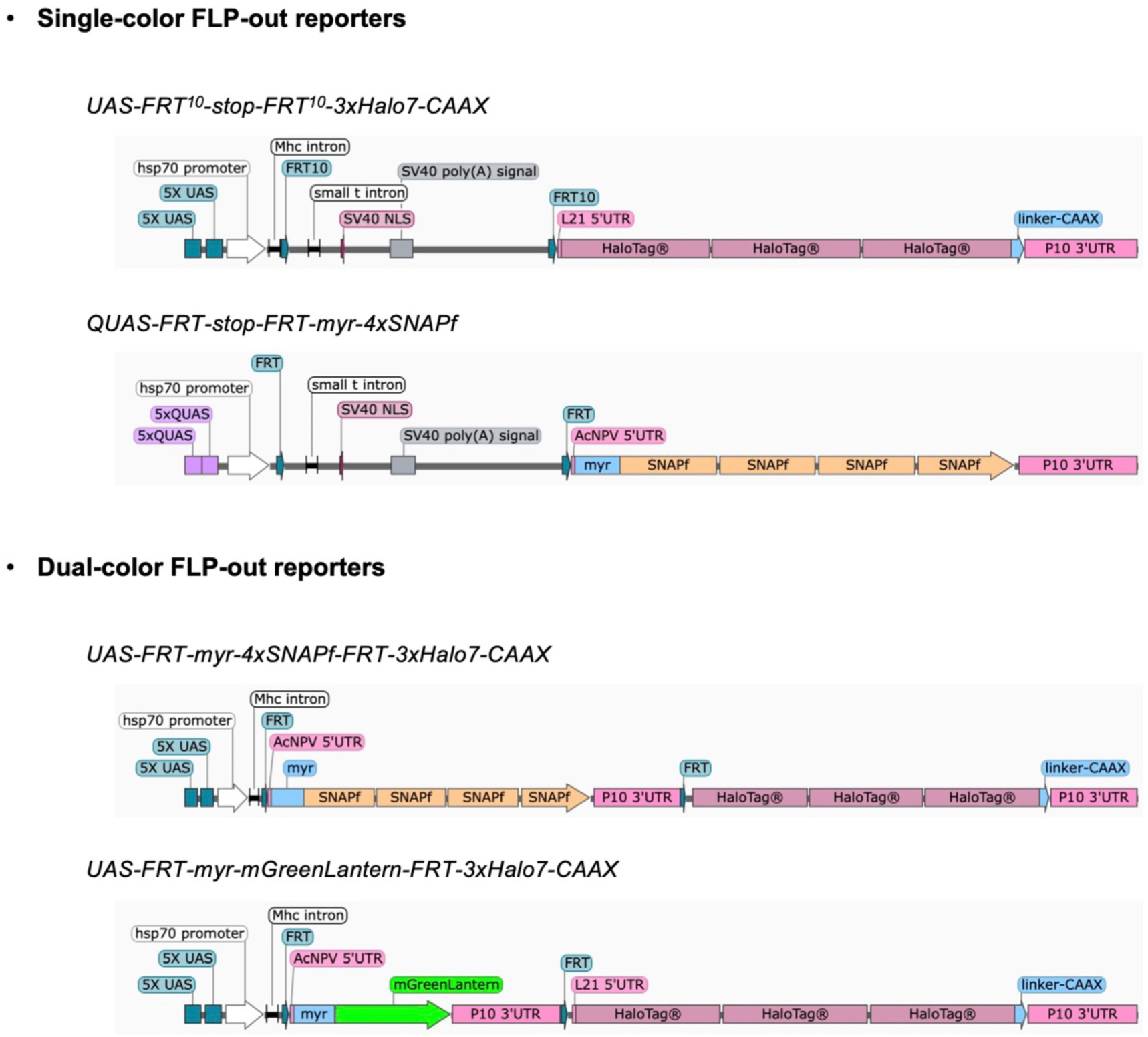
Design of single- and dual-color FLP-out reporters. Images, created with SnapGene, show four newly generated *Q/UAS-*based single- and dual-color FLP-out reporters.

**Figure 3 – figure supplement 1.**
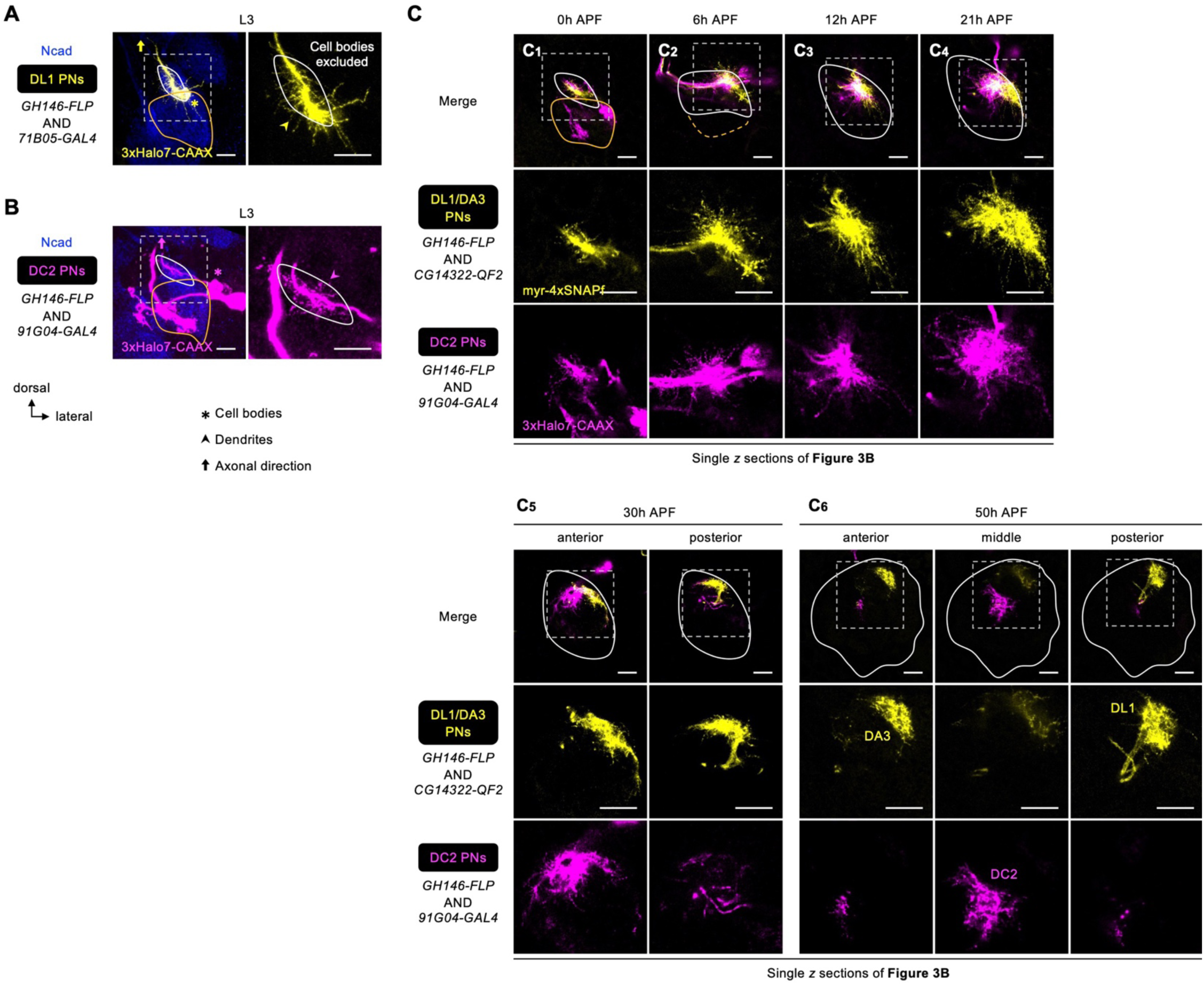
Dendrite development of early larval-born PNs. **(A)** Dendritic extension of DL1 PNs (*71B05+*; labeled in yellow) across the developing antennal lobe at the wandering third instar larval stage (L3). Zoom-in of the dashed box shown on the right. **(B)** Dendritic extension of DC2 PNs (*91G04+*; labeled in yellow) across the developing antennal lobe at L3. Zoom-in of the dashed box shown on the right. **(C)** Single *z* sections of Figure 3B showing dendrite development of DL1/DA3 adPNs (*CG14322+*; labeled in yellow) and DC2 adPNs (*91G04+*; labeled in magenta). See Figure 1 legend for common notations.

**Figure 3 – figure supplement 2.**
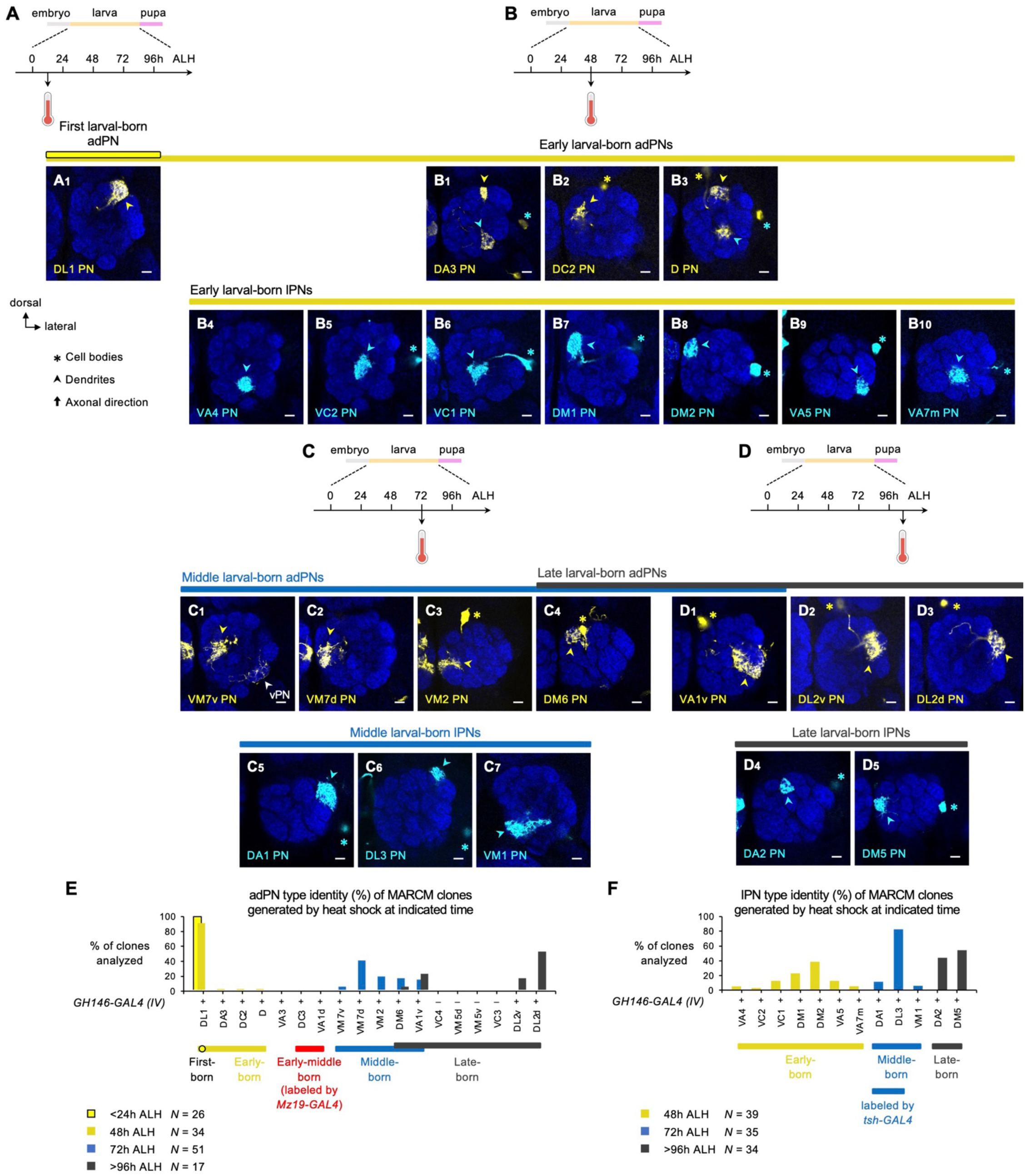
MARCM-labeled single-cell PNs of indicated lineages in adult brains. **(A)** MARCM clone of DL1 PN (in yellow) generated by heat shock at <24h ALH. **(B)** MARCM clones of early larval-born PNs (**B_1–3_**: adPNs in yellow; **B_4–10_**: lPNs in cyan) generated by heat shock at 48h ALH. In **B_1_**, single-cell clone of the adPN lineage and that of the lPN lineage, corresponding to DA3 PN (yellow arrowhead) and VA5 PN (cyan asterisk and arrowhead), were simultaneously generated. In **B_3_**, single-cell adPN and lPN, corresponding to D PN (yellow asterisk and arrowhead) and VA7m PN (cyan asterisk and arrowhead), were simultaneously generated. **(C)** MARCM clones of middle larval-born PNs (**C_1–4_ and D_1_**: adPNs in yellow; **C_5–7_**: lPNs in cyan) generated by heat shock at 72h ALH. In **C_1_**, white arrowhead mark processes of vPN clone that do not belong to VM7v PN. **(D)** MARCM clones of late larval-born PNs (**C_4_ and D_1–3_**: adPNs in yellow; **D_4–5_**: lPNs in cyan) generated by heat shock at >96h ALH. **(E)** Percentage bar graph showing the adPN type identity of MARCM clones generated by heat shock at indicated times. Sample size *N* indicates the number of clones analyzed. **(F)** Percentage bar graph showing the lPN type identity of MARCM clones generated by heat shock at indicated times. Sample size *N* indicates the number of clones analyzed. See Figure 1 legend for common notations.

**Figure 3 – figure supplement 3.**
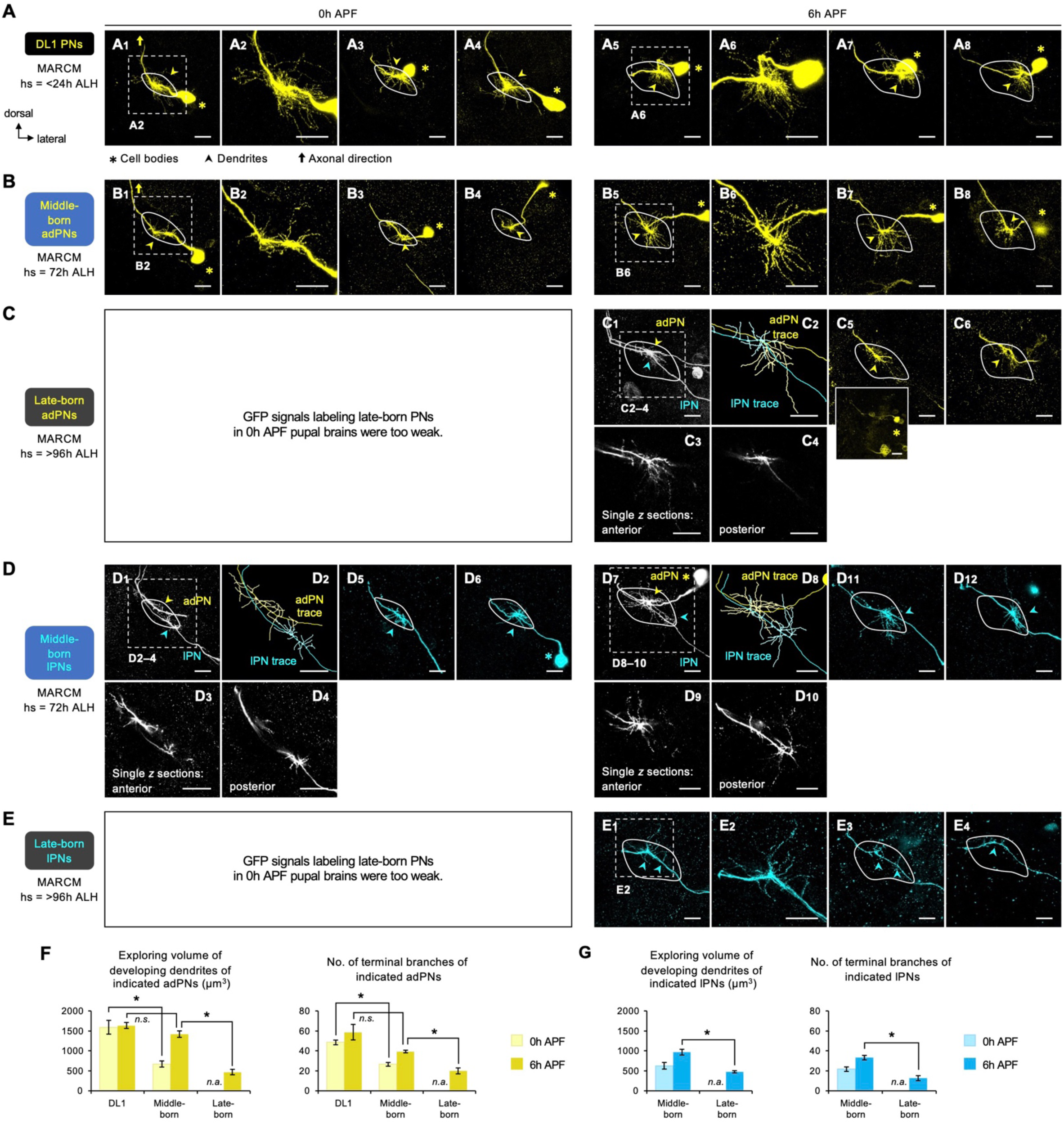
Dendrite development of DL1, middle larval-born, and late larval-born PNs at early stages. Adult-specific antennal lobes (revealed by high Ncad staining; not shown) are outlined by white solid line. **(A)** MARCM clones of DL1 PNs (in yellow), generated by heat shock at <24h ALH, in 0h APF (**A_1–4_**) and 6h APF (**A_5–8_**) pupal brains. **(B)** MARCM clones of middle larval-born adPNs (in yellow), generated by heat shock at 72h ALH, in 0h APF (**B_1–4_**) and 6h APF (**B_5–8_**) pupal brains. **(C)** MARCM clones of late larval-born adPNs (in yellow), generated by heat shock at >96h ALH, in 6h APF pupal brains. In **C_1_**, single-cell MARCM clones of adPN and lPN lineages were simultaneously labeled. **C_2_** shows neurite tracing of adPN (in yellow) and lPN (in cyan) in **C_1_**. Single *z* sections of **C_1_** are shown in **C_3–4_**. Small inset below **C_5_** reveals the cell body position. **_(D)_** MARCM clones of middle larval-born lPNs (in cyan), generated by heat shock at 48h ALH, in 0h APF (**D_1–6_**) and 6h APF (**D_7–12_**) pupal brains. In **D_1–4_** and **D_7–10_**, single-cell adPN and lPN were simultaneously labeled. **D_2_** and **D_8_** shows neurite tracing of **D_1_** and **D_7_**, respectively (adPN in yellow; lPN in cyan). Single *z* sections of **D_1_** are shown in **D_3–4_**, and those of **D_7_** are shown in **D_9–10_**. **(E)** MARCM clones of late larval-born lPNs (in cyan), generated by heat shock at >96h ALH, in 6h APF pupal brains. **(F–G)** Quantification of exploring volume of developing dendrites of indicated PNs (**F:** adPNs; **G:** lPNs) at 0h and 6h APF (left). Quantification of number of terminal branches of indicated PNs (**F:** adPNs; **G:** lPNs) at 0h and 6h APF (right). Error bars, SEM; *t* test; *\**, *p* < 0.05; *n.s.*, *p* ≥ 0.05. SEM, standard error of the mean; *n.s.*, not significant; *n.a.*, not applicable.

**Figure 4 – figure supplement 1.**
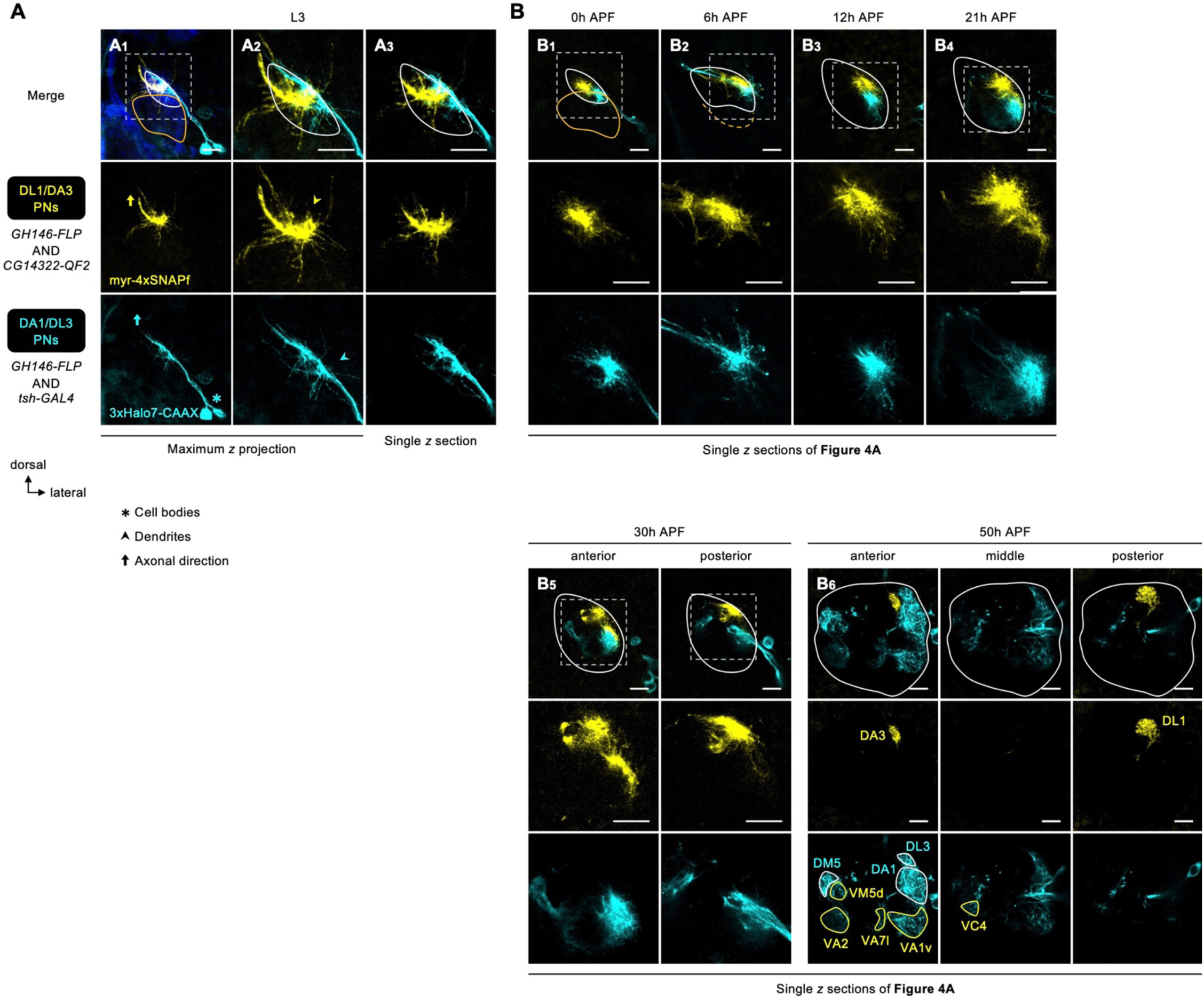
Dendrite development of DL1/DA3 and DA1/DL3 PNs. **(A)** Dendritic extension of DL1/DA3 PNs (*CG14322+*; labeled in yellow) and DA1/DL3 PNs (*tsh+*; labeled in cyan) across the developing antenna lobe at the wandering third instar larval stage (L3). **(B)** Single *z* sections of Figure 4A showing dendrite development of DL1/DA3 PNs (*CG14322+*; labeled in yellow) and DA1/DL3 PNs (*tsh+*; labeled in cyan). In **B_6_**, glomeruli innervated by *tsh+* adPN and *tsh*+ lPN dendrites are outlined in yellow and cyan, respectively. See Figure 1 legend for common notations.

**Figure 4 – figure supplement 2.**
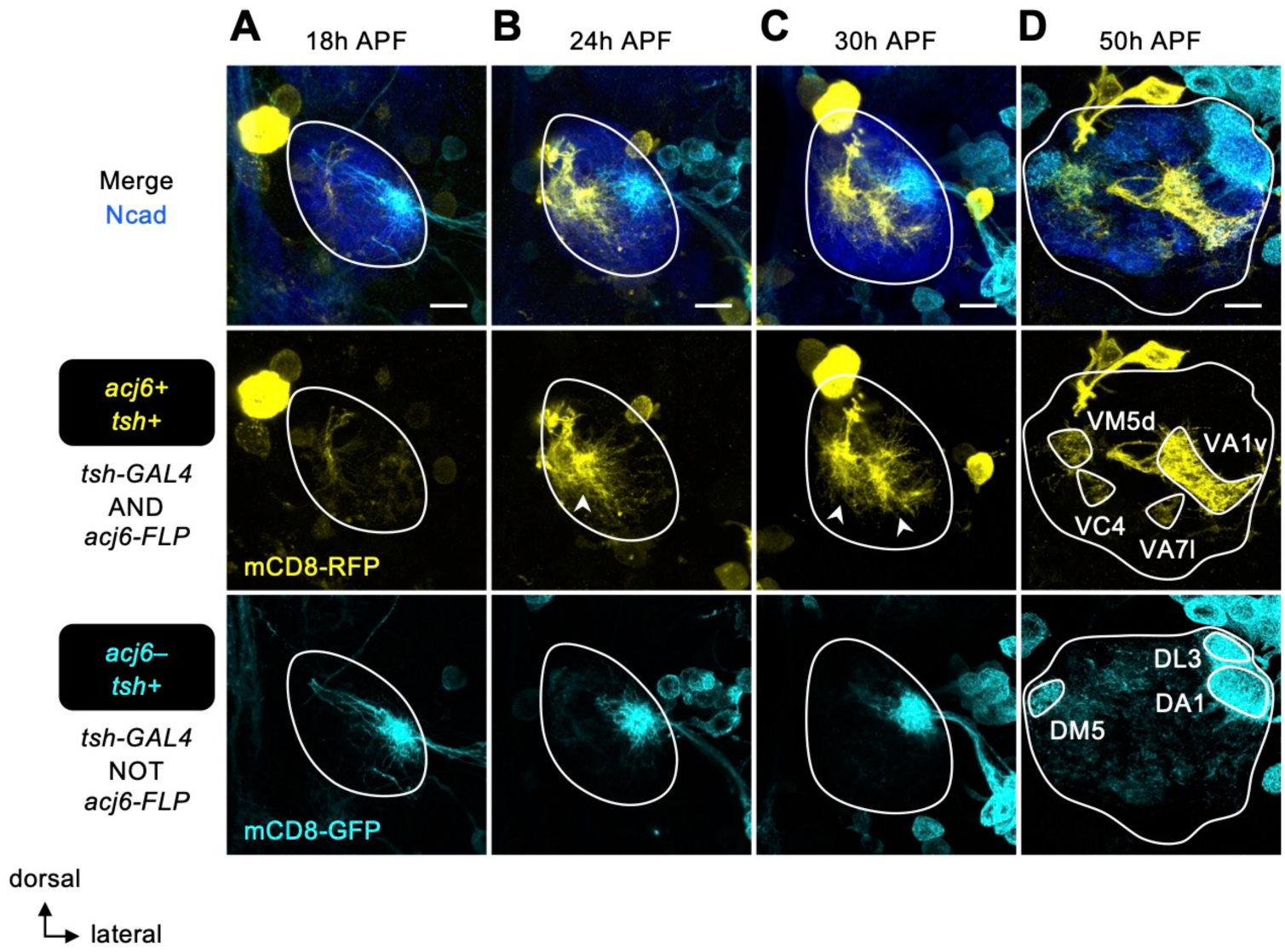
Expression patterns of *tsh* in the developing antennal lobe during mid-pupal stages. **(A)** At 18h APF, *tsh* is only expressed in PNs of the lateral linage (*acj6–*; in cyan). **(B–D)** From 24h APF onwards, *tsh* is expressed in some adPNs (*acj6+*; in yellow), consistent with the transcriptome data (Figure 2B, Figure 2 **– figure supplement 1C–E**). *tsh-GAL4* seems to weakly label local interneurons at 50h APF (*acj6–*; in cyan). See Figure 1 legend for common notations.

**Figure 5 – figure supplement 1.**
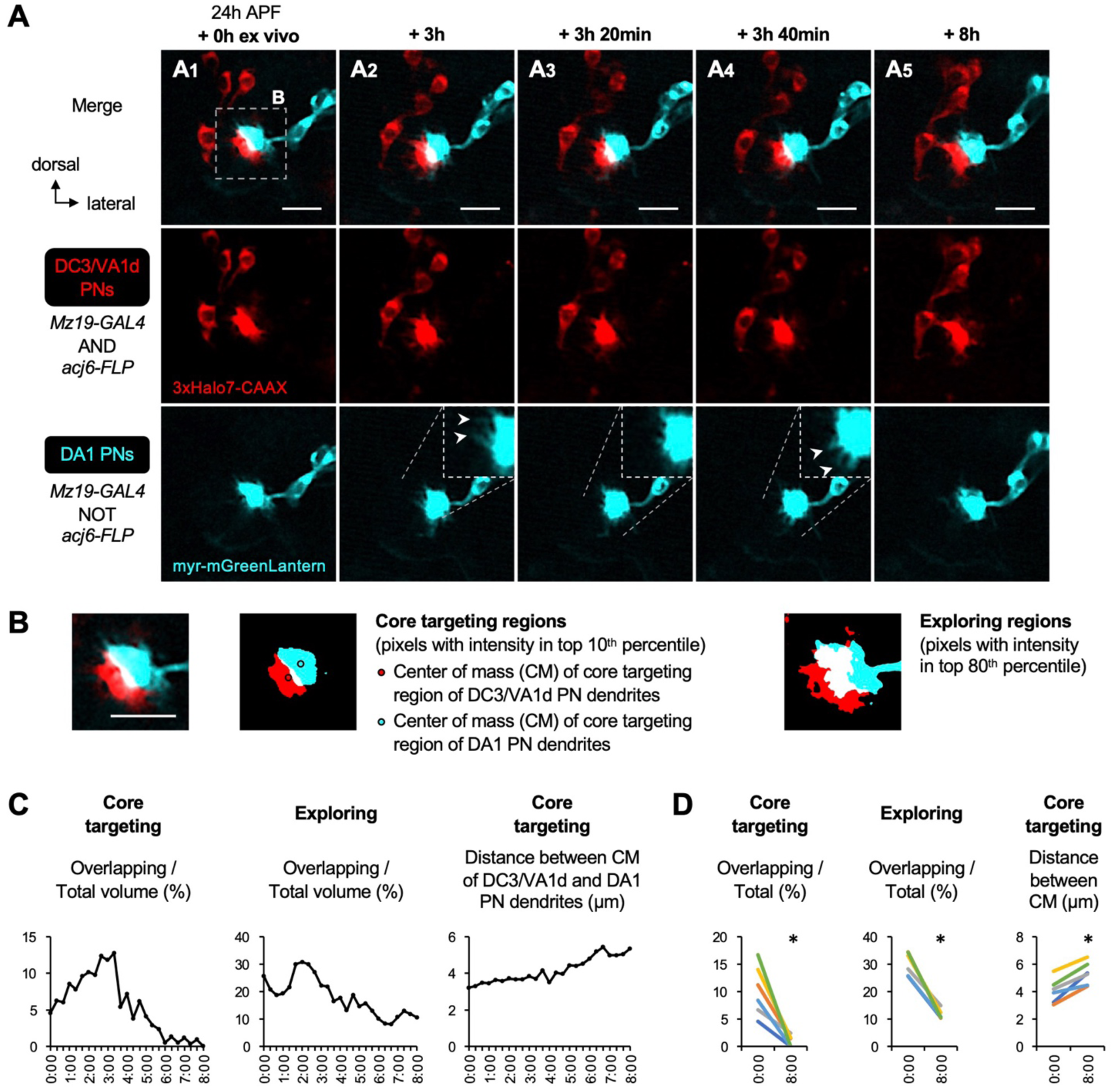
Dendritic segregation of DC3/VA1d adPNs and DA1 lPNs targeting neighboring proto-glomeruli. **(A)** Two-photon time-lapse imaging of DC3/VA1d adPN (*Mz19+ acj6*+; labeled in red) and DA1 lPN (*Mz19+ acj6*–; labeled in red) dendrites in pupal brain dissected at 24h APF and cultured for 8h *ex vivo*. Insets in **A_2–4_** show the zoom-in. Arrowheads in **A_2–4_** indicate the disappearance (compare **A_2_** with **A_3_**) and extension (compare **A_4_** with **A_3_**) of dendrites. **(B)** Core targeting region of PN dendrites is defined using pixels with intensity in the top 10^th^ percentile. Red and cyan circles mark the centers of mass of the core targeting regions of DC3/VA1d and DA1 PN dendrites, respectively. Exploring region of PN dendrites is defined using pixels with intensity in the top 80^th^ percentile. **(C) Left:** Ratio of overlapping to total core targeting volume (in percentage) across the 8-h imaging period. **Middle:** Ratio of overlapping to total exploring volume (in percentage) across the 8-h imaging period. **Right:** Distance between centers of mass of DC3/VA1d and DA1 core targeting regions across the 8-h imaging period. Sample size *N* = 1. Timestamp 00:00 refers to HH:mm; H, hour; m, minute. **(D) Left:** Ratio of overlapping to total core targeting volume (in percentage) at 0h and 8h *ex vivo*. **Middle:** Ratio of overlapping to total exploring volume (in percentage) at 0h and 8h *ex vivo*. **Right:** Distance between centers of mass of DC3/VA1d and DA1 core targeting regions at 0h and 8h *ex vivo*. Sample size *N* = 6. Error bars, standard error of the mean; *t* test; *\**, *p* < 0.05. Timestamp 00:00 refers to HH:mm; H, hour; m, minute. **Figure 5 – video 1. Two-photon time-lapse imaging of PN development.** See Figure 5E for details. Timestamp 00:00:00 refers to HH:mm:ss; H, hour; m, minute; s, second. **Figure 5 – video 2. Two-photon time-lapse imaging of PN dendritic segregation.** See Figure 5 **– figure supplement 1** for details. Timestamp 00:00:00 refers to HH:mm:ss; H, hour; m, minute; s, second.

**Figure 6 – figure supplement 1.**
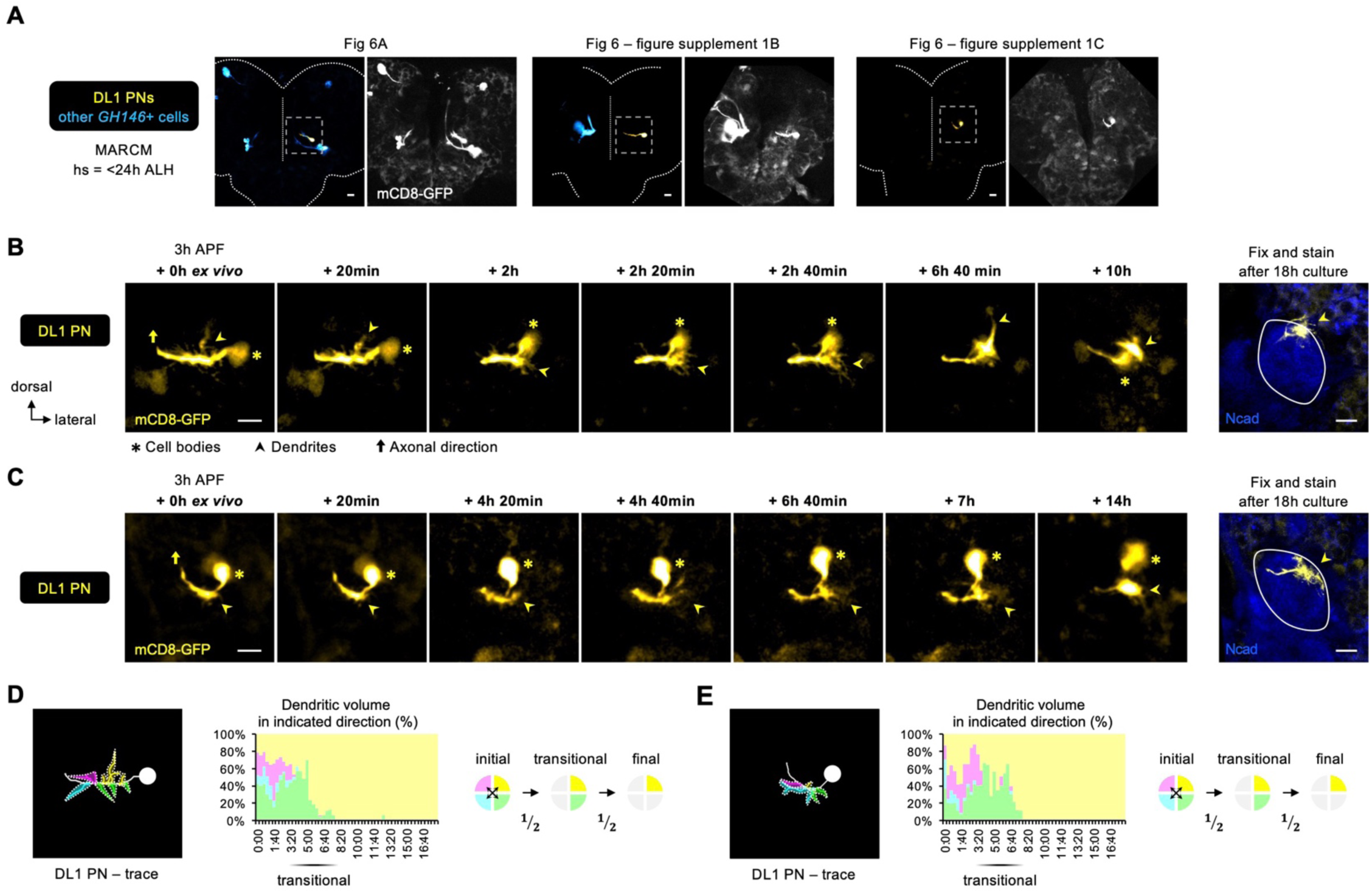
Two-photon time-lapse imaging of DL1 PNs. **(A)** Two-photon images of MARCM-labeled DL1 PNs (pseudo-colored in yellow) and other *GH146*+ cells (pseudo-colored in blue) in pupal brains dissected at 3h APF. Zoom-in time-lapse images of the dashed boxes are shown in Figure 6A, Figure 6 **– figure supplement 1B, C**. Background signals (in grey) are used to discern the orientation of DL1 PN in the brain. **(B, C)** Two-photon time-lapse imaging of MARCM-labeled DL1 PNs (pseudo-colored in yellow) in additional pupal brains dissected at 3h APF and cultured for 18h *ex vivo.* After culture, explant was fixed and immune-stained for N-Cadherin (Ncad; in blue) to outline the developing antennal lobe. **(D, E)** Neurite tracing of DL1 PN (Figure 6 **– figure supplement 1B, C)** at the beginning of live imaging (3h APF + 0h *ex vivo*) and quantification of the percentage of dendritic volume in indicated direction during the time-lapse imaging period. During the transitional period, dendrites are only found in the dorsolateral and ventrolateral directions. Timestamp 00:00 refers to HH:mm; H, hour; m, minute. **Figure 6 – video 1. Two-photon time-lapse imaging of DL1 PN dendrites.** See Figure 6A for details. Timestamp 00:00:00 refers to HH:mm:ss; H, hour; m, minute; s, second.

**Figure 7 – figure supplement 1.**
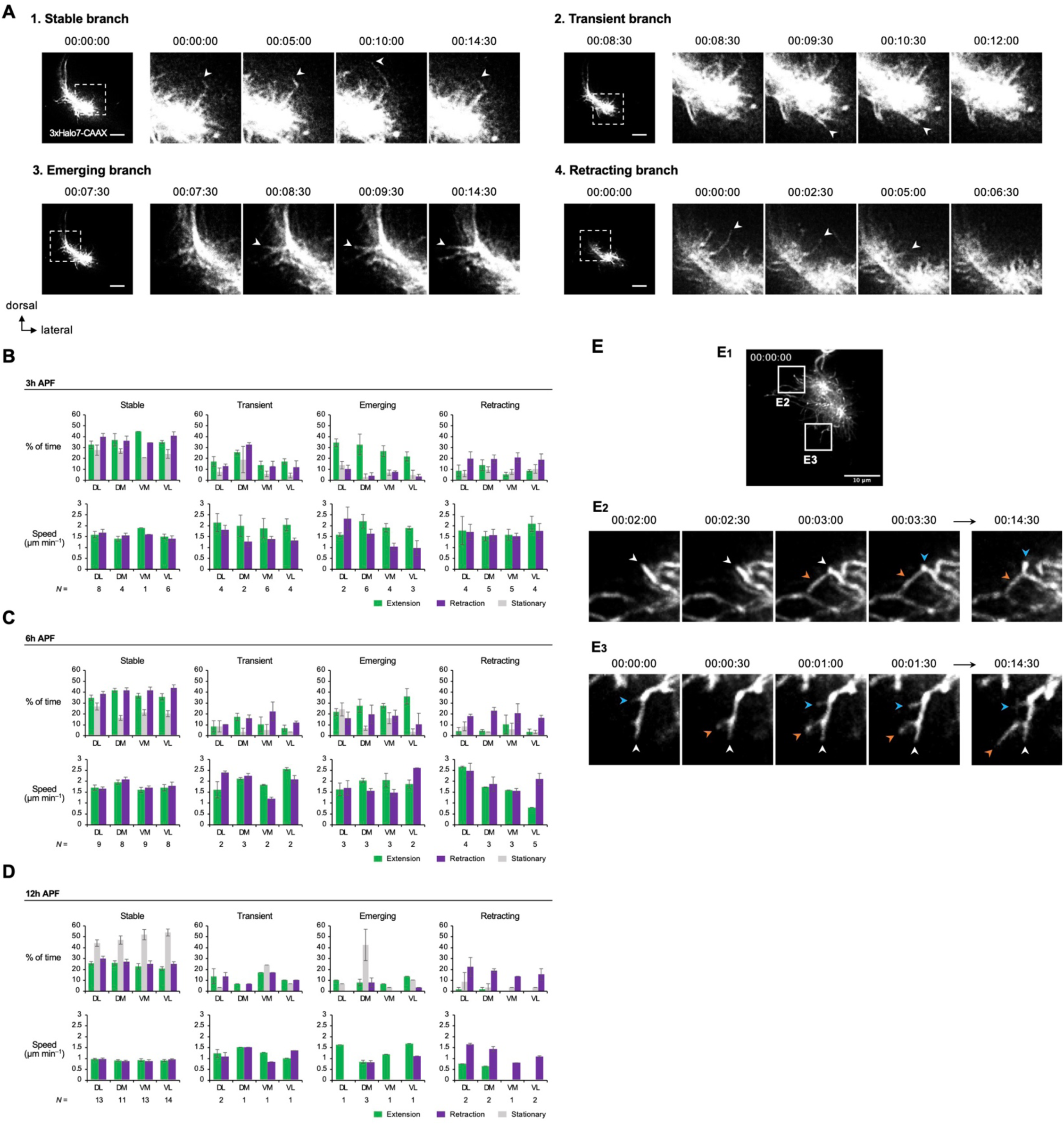
Analyses of DL1 PN dendritic branches captured by AO-LLSM imaging. **(A)** Categorization of branches into **(1)** stable, **(2)** transient, **(3)** emerging, and **(4)** retracting branches. Representative branches of each category are shown. Timestamp 00:00:00 refers to HH:mm:ss; H, hour; m, minute; s, second. **(B–D) Top:** Quantification of the percentage of time given branches spent on extending, retracting, and being stationary. **Bottom:** Extension/retraction speeds of DL1 PN stable, transient, emerging, and retracting branches in indicated directions at 3h APF **(B)**, 6h APF **(C)**, and 12h APF **(D)**. Data are analyzed using DL1 PNs in Figure 7A**–C**. *N* indicates branch number. **(E)** Time-lapse AO-LLSM imaging of 12h APF DL1 PNs (**E_1_**; Figure 7C) reveals terminal branch arborization. **E_2_** and **E_3_** are selected time-lapse images of zoom-in of indicated boxes in **E_1_**. White arrowheads point to the terminal branch of interest, and colored arrowheads point to secondary branches produced from the branch of interest. Timestamp 00:00:00 refers to HH:mm:ss; H, hour; m, minute; s, second. **Figure 7 – video 1. AO-LLSM time-lapse imaging of 3h DL1 PN dendrites.** See Figure 7A for details. Timestamp 00:00:00 refers to HH:mm:ss; H, hour; m, minute; s, second. **Figure 7 – video 2. AO-LLSM time-lapse imaging of 6h DL1 PN dendrites.** See Figure 7B for details. Timestamp 00:00:00 refers to HH:mm:ss; H, hour; m, minute; s, second. **Figure 7 – video 3. AO-LLSM time-lapse imaging of 12h DL1 PN dendrites.** See Figure 7C for details. Timestamp 00:00:00 refers to HH:mm:ss; H, hour; m, minute; s, second.

**Figure 8 – figure supplement 1.**
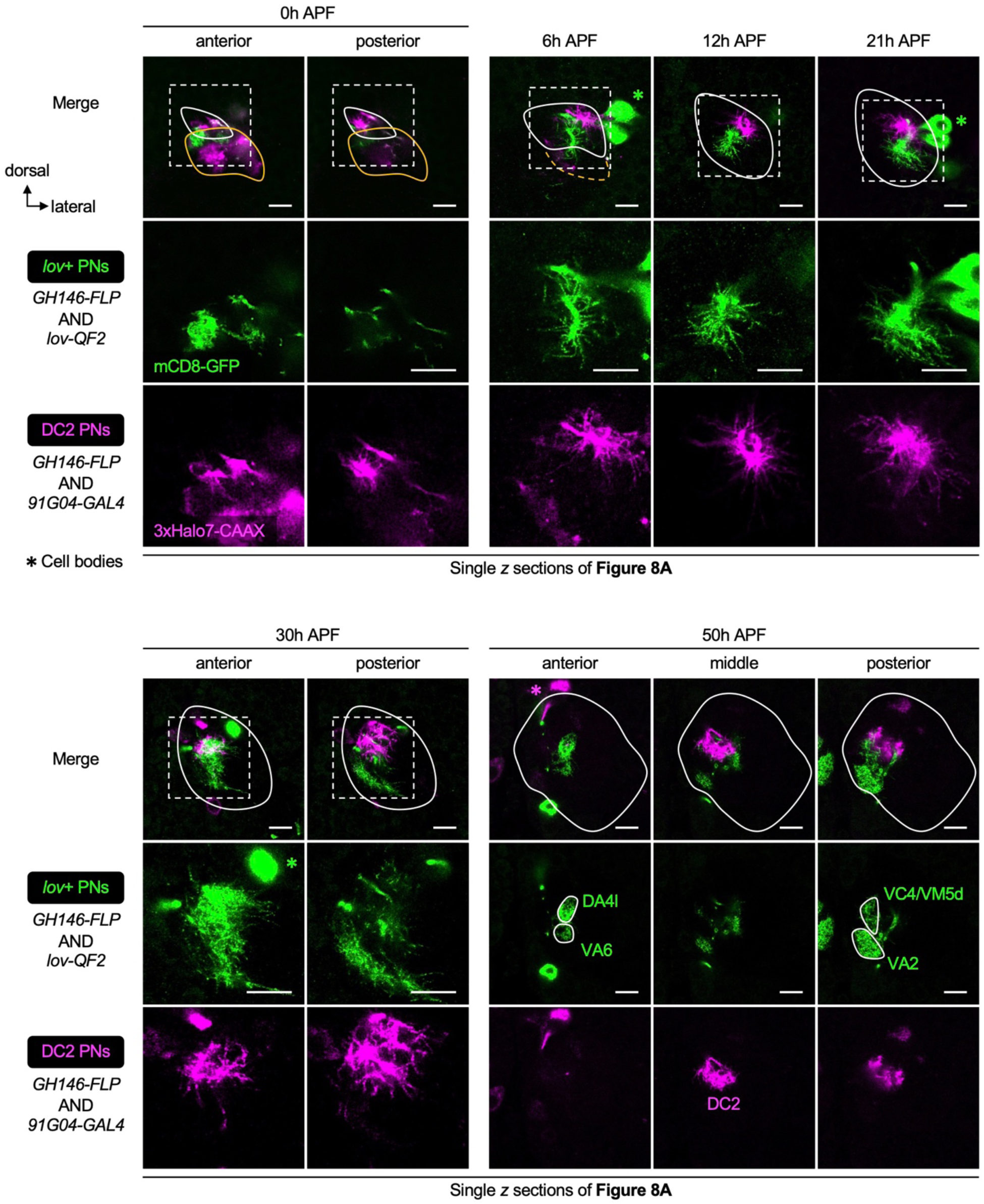
**Dendrite development of lov+ embryonic-born PNs.** Single *z* sections of Figure 8A showing the dendrite development of *lov+* PNs (embryonic-born; labeled in green) and *91G04+* DC2 PNs (larval-born; labeled in magenta).

**Figure 8 – figure supplement 2.**
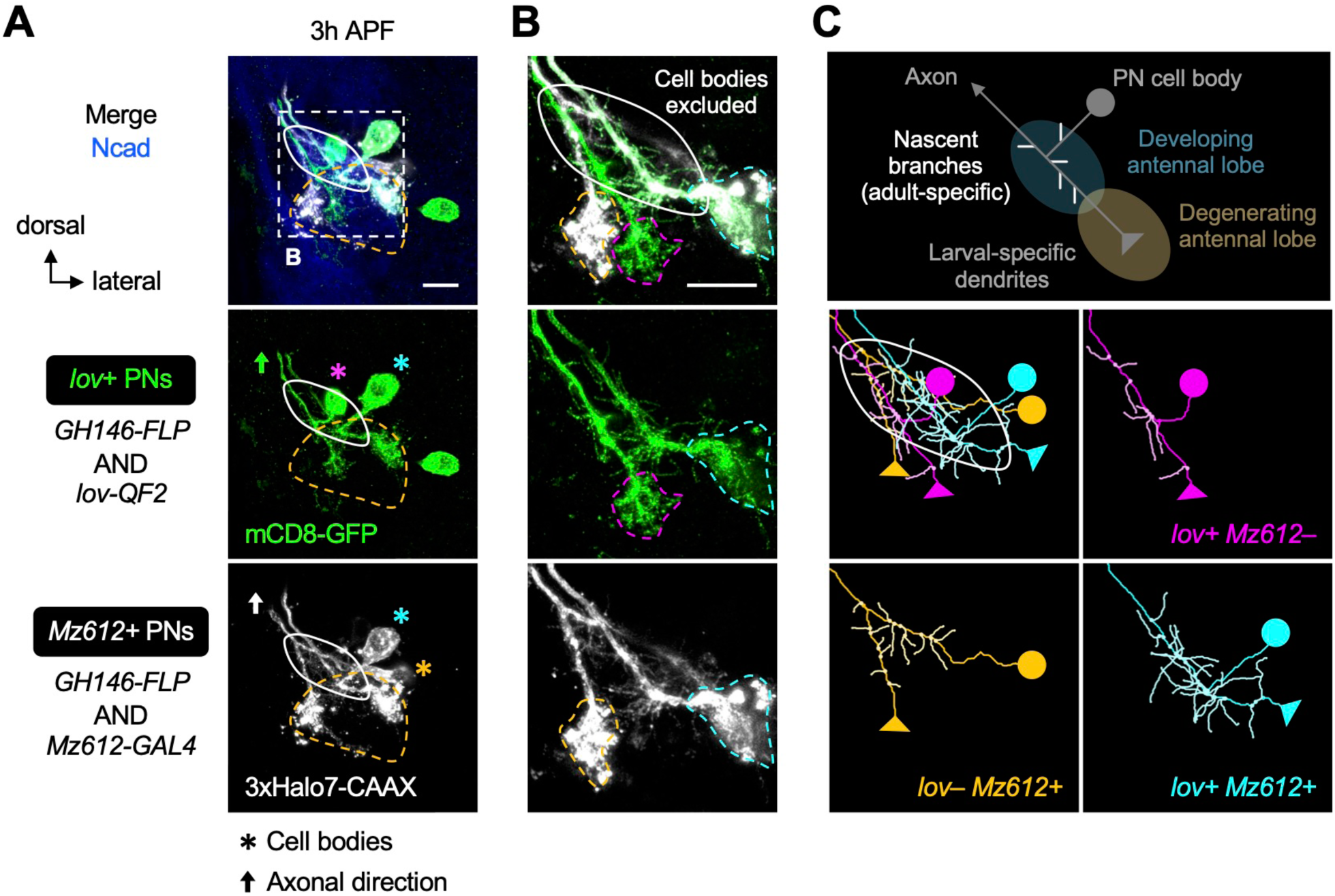
Dendrite re-extension of lov+ and Mz612+ embryonic-born PNs. **(A, B)** Dendritic re-extension of *lov+* (labeled in green) and *Mz612+* (labeled in grey) PNs. **B** is the zoom-in of dashed box in **A**. Magenta, cyan, and orange asterisks in **A** indicate cell bodies of *lov+ Mz612–* PN, *lov+ Mz612*+ PN, and *lov– Mz612*+ PN, respectively. Using the same color code, their larval-specific dendrites are outlined with dashed lines in **B**. **(C)** Schematic of co-existence of larval- and adult-specific dendrites of an embryonic-born PN (top row), and neurite tracing of the three embryonic-born PNs (middle and bottom rows).

**Figure 8 – figure supplement 3.**
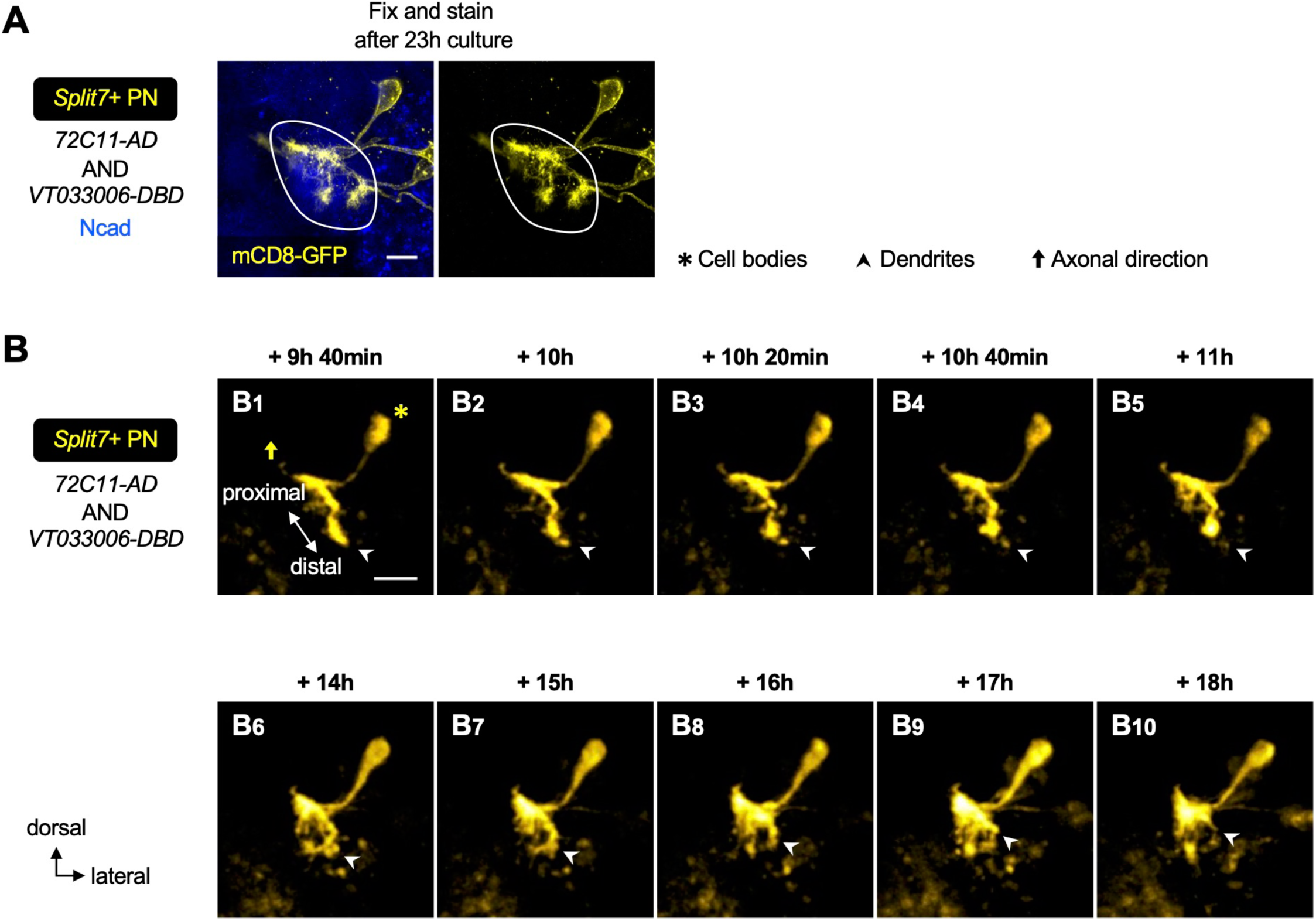
Two-photon time-lapse imaging of *Split7+* PN dendrites. **(A)** Confocal image of *Split7+* PNs (labeled in yellow; Figure 8D) after 23h culture. N-Cadherin (Ncad; in blue) staining outlines the developing antennal lobe. *Split7-GAL4* is expressed in more than one PN type at later stages. **(B)** Two-photon time-lapse imaging of a *Split7+* PN showing distal-to-proximal pruning of larval-specific dendrites.

**Figure 8 – figure supplement 4.**
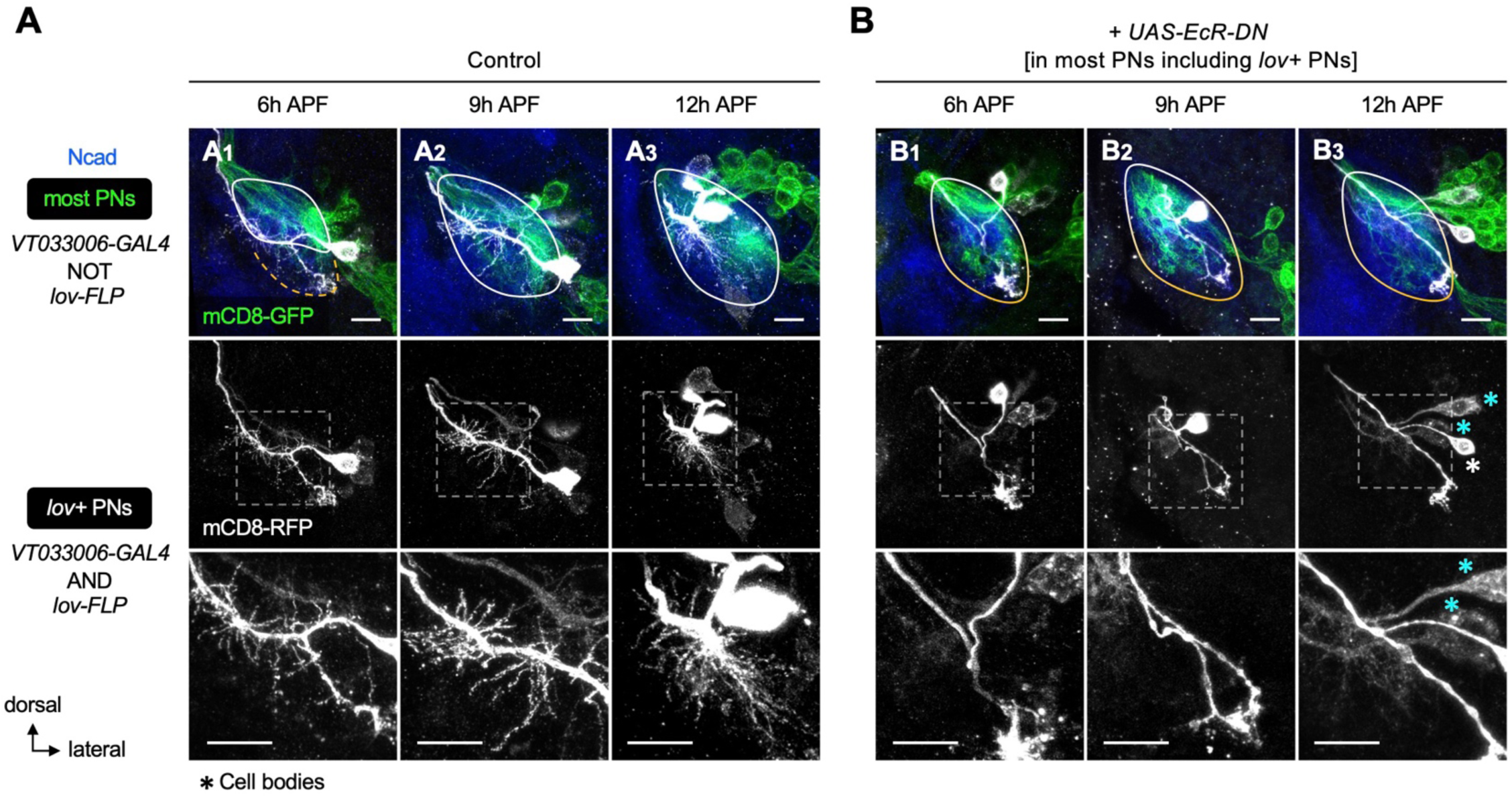
Dual requirement of ecdysone signaling in pruning and re-extension of embryonic-born PN dendrites. **(A)** Normal dendrite development seen in control *lov*+ embryonic-born PNs (*VT033006+ lov+*; labeled in grey). Other PNs (referred to as most PNs; *VT033006+ lov–*) are labeled in green. Bottom row shows zoom-in of the dashed boxes. **(B)** Expression of a dominant negative form of ecdysone receptor (*EcR-DN*) in most PNs including *lov+* PNs suppresses both pruning and re-extension of *lov+* PN dendrites. Similar results were seen in multiple biological samples (*N* ≥ 3 for each stage per genotype). Cyan asterisks in **B_3_** mark *lov+* cells that had weak *VT033006- GAL4* and thereby weak *EcR-DN* expression. These cells appeared to still elaborate dendrites, suggestive of a dose-dependent effect of *EcR-DN*. White asterisk in **B_3_** mark a *lov+* cell with strong *VT033006-GAL4* expression. The presumed fused larval- and adult-specific antennal lobes are outlined with white-orange gradient line. **Figure 8 – vid**eo 1. Two-photon time-lapse imaging of *Split7+* PN dendrites. See Figure 8D for details. Timestamp 00:00:00 refers to HH:mm:ss; H, hour; m, minute; s, second.

